# Therapeutic advantages of combined gene/cell therapy strategies in a murine model of GM2 gangliosidosis

**DOI:** 10.1101/2021.12.22.473777

**Authors:** Davide Sala, Francesca Ornaghi, Francesco Morena, Chiara Argentati, Manuela Valsecchi, Valeria Alberizzi, Roberta Di Guardo, Alessandra Bolino, Massimo Aureli, Sabata Martino, Angela Gritti

## Abstract

The GM2 gangliosidoses Tay-Sachs disease and Sandhoff disease (SD) are respectively caused by mutations in the HEXA and HEXB genes encoding the α and β subunits of β-N-acetylhexosaminidase (Hex). The consequential accumulation of ganglioside in the brain leads to severe and progressive neurological impairment. There are currently no approved therapies to counteract or reverse the effects of GM2 gangliosidosis. Adeno-associated vector (AAV)-based investigational gene therapy (GT) products have raised expectations but come with safety and efficacy issues that need to be addressed. Thus, there is an urgent need to develop novel therapies targeting the CNS and other affected tissues that are appropriately timed to ensure pervasive metabolic correction and counteract disease progression. In this report, we show that the sequential administration of lentiviral vector (LV)-mediated intracerebral (IC) GT and bone marrow transplantation (BMT) in pre-symptomatic SD mice provide a timely and long-lasting source of the Hex enzyme in the central and peripheral nervous systems and peripheral tissues, leading to global rescue of the disease phenotype. Combined therapy showed a clear therapeutic advantage compared to individual treatments in terms of lifespan extension and normalization of the neuroinflammatory and neurodegenerative phenotypes of the SD mice. These benefits correlated with a time-dependent increase in Hex activity and a remarkable reduction in GM2 storage in the brain tissues that single treatments failed to achieve. Our results highlight the complementary and synergic mode of action of LV-mediated IC GT and BMT, clarify the relative contribution of treatments to the therapeutic outcome, and inform on the realistic threshold of enzymatic activity that is required to achieve a significant therapeutic benefit, with important implications for the monitoring and interpretation of ongoing experimental therapies, and for the design of more effective treatment strategies for GM2 gangliosidosis.

## INTRODUCTION

Genetic defects in the β-N-acetylhexosaminidase (Hex) enzyme cause GM2 gangliosidoses, a group of rare neurodegenerative lysosomal storage diseases (LSD). This enzymatic deficiency leads to the accumulation of GM2 and other glycosphingolipids in the central nervous system (CNS) and other organs (1). The Hex enzymes are found in three isoenzymes resulting from variations in the association of different α and β subunits (HexA,αβ; HexB, ββ; HexS, αα) encoded by the HEXA (mutated in Tay-Sachs disease, TSD) and HEXB genes (mutated in Sandhoff disease, SD). The infantile forms of GM2 gangliosidosis are the most clinically common types and are extremely severe, but attenuated juvenile and late-onset forms also occur (2–4).

There are currently no treatments for GM2 gangliosidosis. Enzyme-replacement therapy (5, 6), substrate-reduction therapy (7, 8), and pharmacological chaperones (9, 10) provide partial benefits in cases of the juvenile/late-onset forms but are ineffective in infantile patients (11–16). Bone marrow transplant (BMT) ameliorates neuroinflammation and prolongs the lifespan of Sandhoff mice (SD; Hexb^−/−^), which develop a progressive and severe neurological phenotype resulting in death at around 4 months of age (17). However, BMT fails to supply therapeutic levels of Hex to CNS tissues, which retain a substantial GM2 storage burden (18, 19). In line with these observations, BMT provides minimal benefit to either TSD or SD patients (20, 21).

Gene therapy (GT) holds promise as a means to address TSD and SD pathology. In vivo intracerebral GT (IC GT) delivering functional Hex enzymes by means of adeno-associated viral (AAV) vectors has benefited SD mice and cats (22–26) and is currently under clinical testing for infantile patients (NCT04798235, NCT04669535). Despite this, pre-clinical studies and clinical observations have highlighted several major issues associated with this strategy, i.e., the necessity to co-deliver the α and β subunits at the appropriate ratio to achieve a fully functional enzyme, the toxicity associated with Hex overexpression in brain cells upon IC GT in non-human primates (27), the immunogenicity of AAVs that require some immunosuppressive treatment administered transiently or long-term to support durable transgene expression (28, 29), and the potential off-target toxicity due to AAV leakiness (30), which demand long-term efficacy and safety evaluations. Lentiviral vectors (LV) transduce neuronal and glial cells with high efficiency and display little dispersion and low immunogenicity upon IC delivery (31, 32), thus representing a complementary/alternative GT platform to target the CNS. The results of the first clinical trial using a LV-based GT vector to treat Parkinson’s disease patients showed a favorable safety profile and indication of efficacy (33, 34). We have previously shown the safety and efficacy of a LV-based IC GT platform to provide stable levels of therapeutic lysosomal enzymes in murine and non-human primate models of leukodystrophies (31, 32, 35). In addition, we have generated mono- and bicistronic LVs driving the expression of Hex genes in neural and hematopoietic cells (36) thus supporting the rationale of testing these LVs in GT platforms to treat GM2 gangliosidosis.

Low levels of GM2 storage may affect developing neurons and glial cells, greatly restricting the time window available for effective therapeutic interventions. Indeed, neurodevelopmental defects have been described in GM2 gangliosidosis (37–39) as well as other infantile neurodegenerative LSDs, e.g., Mucopolysaccharidosis type 1 (MPSI) and Nieman-Pick (40, 41), and globoid cell leukodystrophy (GLD) (42, 43). Further to the timing of the intervention, the threshold concentration of enzyme needed for therapeutic effect is a central issue for consideration. It is widely assumed that 10-15% of normal enzymatic activity is sufficient to hydrolyze excess substrates and prevent disease manifestations in various LSDs, including GM2 gangliosidosis (2, 44, 45). However, whether a treatment ensures these enzymatic levels are met on a per cell basis in animal models and patients’ needs to be clarified. In addition, the early neuroinflammation and myelin dysfunction described in GM2 gangliosidosis animal models and patients (19, 22, 46, 47) suggest that a therapy able to limit and prevent tissue damage and promote repair, in addition to restoring an endogenous source of functional enzyme, is required. Finally, despite TSD and SD primarily affect the CNS, approaches that provide additional enzymatic correction and neuroprotection to the PNS and peripheral organs would likely enhance the therapeutic benefits. To this end, a variable synergy in terms of clearance of storage, survival, and functional rescue has been described when combining oral glycosphingolipid biosynthesis inhibitors with neural stem cell transplant (NSCT) (48, 49) or BMT (50). We have previously shown that lentiviral vector (LV)-mediated neonatal NSCT or IC GT synergizes with BMT, providing remarkable health benefits to a severe murine model of GLD (51). The results of our study indicated that the early availability of functional enzyme promoted by NSCT and IC GT is instrumental to enhancing the long-term advantages of BMT. These findings provide a strong rationale to evaluate the therapeutic benefits of combined strategies based on IC GT and BMT in GM2 gangliosidosis diseases, which share with GLD the problems of early onset, rapid progression, severe CNS damage, and limitations to the benefits achieved after experimental therapies are applied individually.

To address the safety and efficacy issues of GT approaches currently being developed (such as *in vivo* AAV GT) we aimed to evaluate the potential of using the LV-mediated GT platform to deliver Hex genes to the CNS and demonstrate the potential of combining this strategy with BMT to prevent/delay disease onset and progression, prolong lifespan, and correct pathological hallmarks of SD mice. We demonstrate that the sequential administration of LV-mediated IC GT and BMT in pre-symptomatic SD mice provides a timely and long-lasting source of Hex as well as conferring neuroprotection/immunomodulation in the CNS, PNS, and periphery. Combined therapy comes with remarkable advantages compared to individual treatments in terms of extending the lifespan of SD mice and global rescue of the disease phenotype. These benefits rely on the complementary modality of the treatments’ mechanisms of action and correlate with increased enzymatic activity and a significant reduction in GM2 storage in CNS tissues.

This work demonstrated the suitability of using an LV-mediated GT platform to deliver Hex genes to the CNS and the overall therapeutic potential of the proposed combined strategy. Additionally, the findings clarify the relative contribution of treatments to the therapeutic outcome, and inform on the realistic threshold of enzymatic activity that is required to achieve a significant therapeutic benefit, with important implications for the monitoring and interpretation of ongoing experimental therapies, and for the design of more effective treatment strategies for GM2 gangliosidosis.

## RESULTS

### Experimental plan

In this study, we took advantage of optimized experimental protocols for LV-mediated IC GT and BMT (31, 32, 51, 52), and the experimental plan is summarized in **Figure 1**. We used LVs to drive the expression of the murine α and β Hex subunits under the control of the human phosphoglycerate kinase (PGK) promoter (LV.hexB and LV.hexA) (53). Pilot experiments showed there was local rescue of enzyme activity and the isoenzyme composition of the injected hemisphere but little enzymatic activity in the contralateral hemisphere upon unilateral LV injection (**Supplementary Figure 1A-C**). Thus, we performed bilateral co-injections of purified LVs into the external capsule (EC) of postnatal day 2 SD mice (Hexb^-/-^)(LV.hexB+LV.hexA; 2 × 10^6^ TU/µl of each vector; 4 µl in total). In the combined treatment group, LV-injected mice at 60-70 d of age were transplanted with total bone marrow (BM) from transgenic green fluorescent protein (tgGFP)-expressing mice. Untreated (UT), IC GT- and BMT-treated SD mice, UT wildtype (WT; Hexb^+/+^), and heterozygous (Het; Hexb^+/-^) age-matched littermates were used as controls. Mice were analyzed at different timepoints during disease progression and after treatment to assess the (i) engraftment of transplanted cells and brain cell transduction upon IC GT; (ii) Hex activity in CNS, PNS, and peripheral organs; and (iii) progression of disease-associated pathological features (lysosomal expansion, glycolipid storage, neuroinflammation, neurodegeneration, and myelin damage) and their treatment-associated rescue by means of a thorough molecular, biochemical, and morphological evaluation.

**Figure 1.**
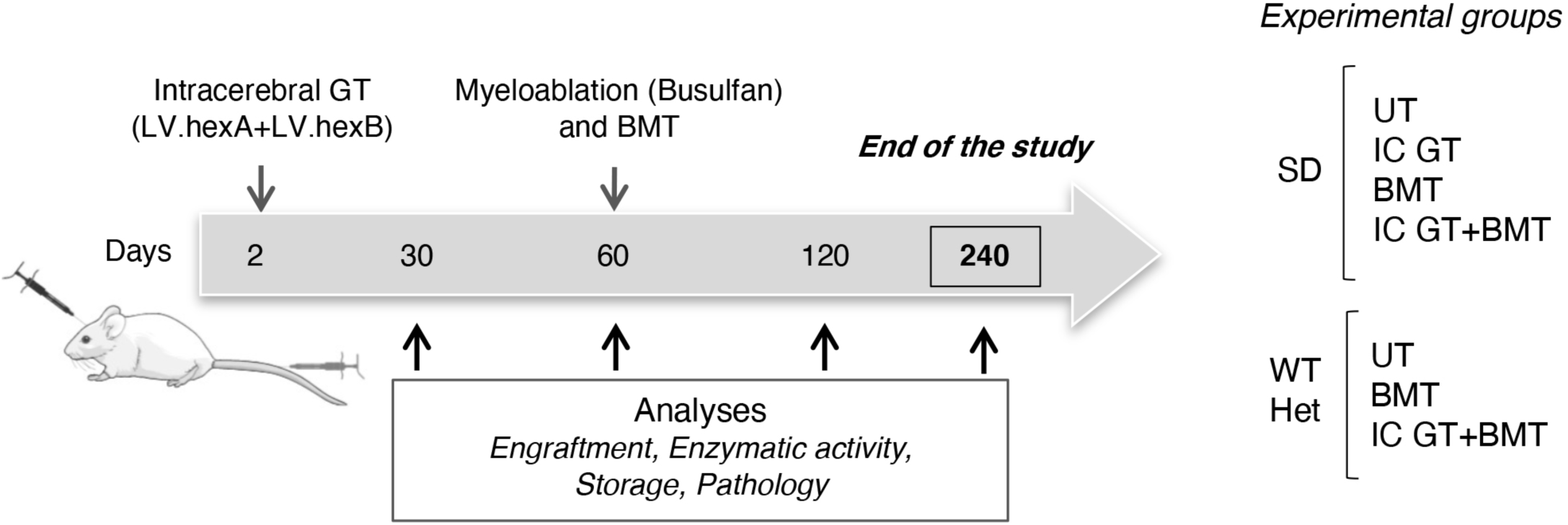
Experimental plan. LVs (LV.hexa+LV.hexb) were co-injected bilaterally into the external capsule of neonatal mice at 2 d. Transplantation of total bone marrow isolated from tgGFP WT donors was performed at 60 d following busulfan treatment (myeloablation). Untreated (UT) and/or treated SD mice (Hexb^-/-^) were analyzed during disease progression (30 d, 60 d, asymptomatic; 120 d, terminal stage of disease) and after treatment (120 d; 240 d, end of the study). Intracerebral gene therapy (IC GT), bone marrow-transplant (BMT), and combined (IC GT+BMT) treatments were applied to SD mice; WT (Hexb^+/+^) and heterozygous (Het; Hexb^+/-^) littermates.

### Synergy of treatments for increasing the lifespan of SD mice

The Kaplan–Meier survival curves for all treatment groups are reported in **Figure 2A**. UT SD mice followed a rapidly progressive neurodegenerative course, starting from around 90 d of age, and showed a median survival of 120 d (100-135 d, n = 93 mice). Single treatments resulted in a modest but significant increase in the median survival, which were 132 d (118-143 d, n = 6) for IC GT and 136 d (122-180 d, n = 13) for BMT. Combination-treated SD mice lived significantly longer than UT SD and SD mice treated with individual approaches, with a mean survival of 210 d (142-246 d; n = 13; with six animals sacrificed at 240 d, the endpoint of the study). Despite the significant delay in the onset of symptoms, both IC GT- and BMT-treated SD mice experienced the rapid and severe disease progression that characterized the UT SD controls, eventually exhibiting muscle wasting, rigidity, and a nearly complete inability to ambulate. In contrast, the combined therapy resulted in strikingly improved clinical appearance at a time when the SD mice treated with BMT or IC GT alone, as well as UT controls, were severely affected with disease manifestations. Indeed, all the combination-treated SD mice (including the three animals sacrificed at 240 d) displayed only mild tremor and ataxia, maintained walking and explorative ability (see **Supplementary videos 1-3**), and were able to feed independently, as well as showing the preservation of normal body weight (**Figure 2B**).

**Figure 2.**
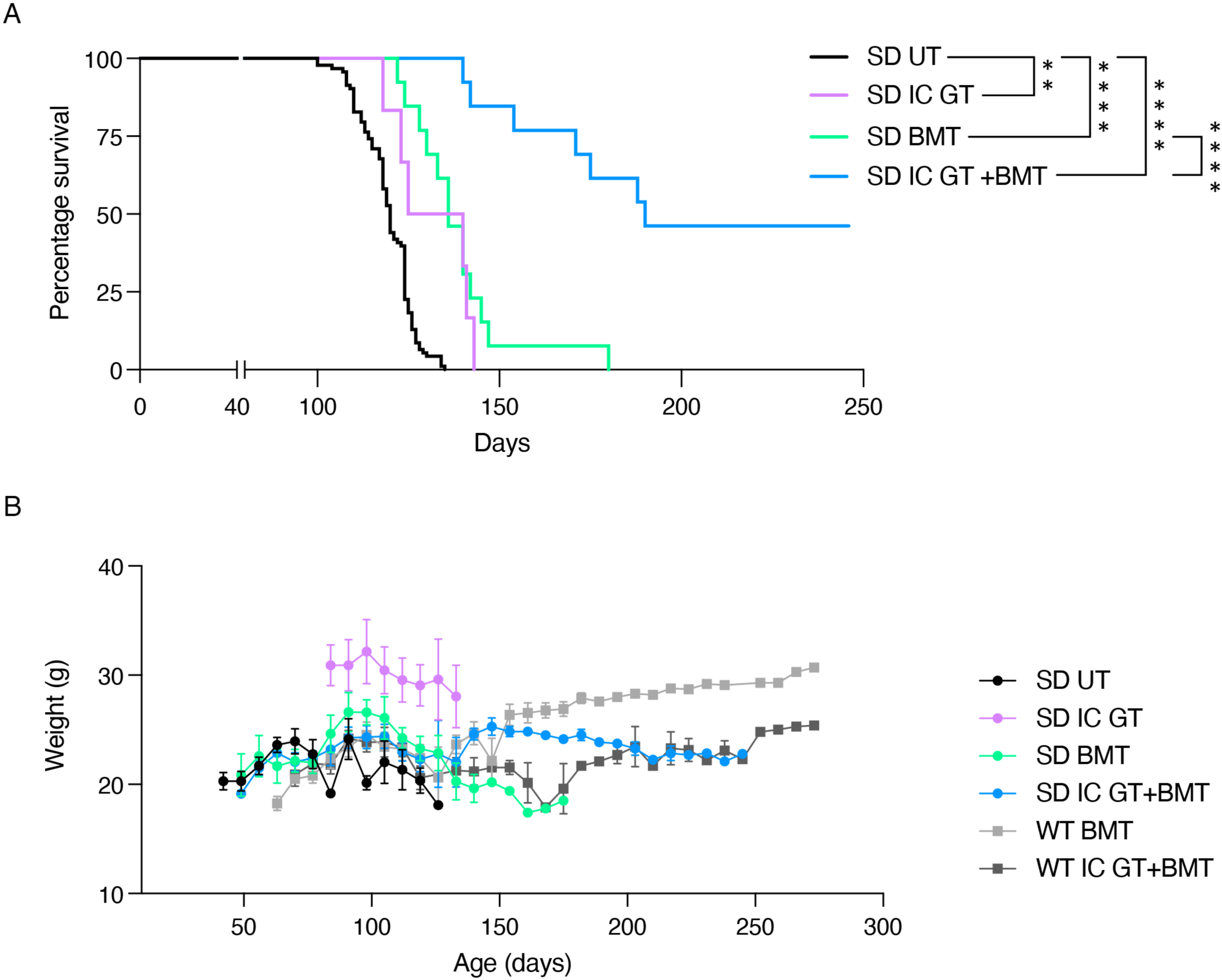
Synergy of treatments in increasing the lifespan and preserving the body weight of SD mice. **(A)** Kaplan–Meier survival curves for the percentage survival of treated and untreated (UT) SD mice. SD UT, n = 93; SD IC GT = 6; SD BMT, n = 13; SD IC GT+BMT, n = 12. Data were analyzed by log-rank (Mantel–Cox) test: ^**^p < 0.005, ^****^p < 0.0001. **(B)** Body weight of treated SD mice and UT littermates was monitored from 50 days of age. BMT-treated WT/Het mice were used as controls to evaluate the effect of busulfan conditioning on body weight. The higher body weight of IC GT-treated mice reflected the absence of myeloablative regimen in this treatment group. Data represent mean ± SEM; n = 5-19 mice/group

### Stable transduction of CNS cells and hematopoietic cell engraftment in combination-treated SD mice

Mice were examined to evaluate the efficacy of neural cell transduction upon IC GT (30 d after LV injection) and the engraftment of BM-derived hematopoietic cells (HCs) in peripheral blood (PB) at 30 d and BM at 60 d after transplantation. In line with our previous studies (51), we detected the robust transduction of endogenous cells and widespread transgene diffusion along white matter tracts upon neonatal IC GT, which we assessed using LV.GFP delivery in a group of control animals (**Supplementary Figure 1D**). Injection of LV.hexA+LV.hexB resulted in a decrease in GM2 storage and a reduction in Cluster of differentiation 68 (CD68)+ macrophages in areas of the brain close to the injection site (**Figure 3A**; 120 d), suggesting the functionality of the transgenic enzyme and the short range of action. The pervasive engraftment of metabolically competent HCs into the CNS tissues of BMT-treated SD mice was expected to enhance the enzymatic supply to therapeutically benefit affected tissues. We applied busulfan as a conditioning regimen capable of ablating the resident hematopoietic niche, as well as brain-resident myeloid precursors, thus favoring the turnover from resident myeloid cells to the donor counterparts (54). Cytofluorimetric analysis showed sustained and stable chimerism in BMT- and combination-treated SD mice (>90% and >60% of GFP+ donor-derived HCs in the PB and BM, respectively; **Figure 3B-C**), with complete reconstitution of white and red blood cell (WBC and RBC) populations and normal WBC composition (**Supplementary Table 1**). Transplanted mice and tgGFP donor mice showed comparable percentages of GFP+ cells and BM cell type compositions (**Figure 3C-D)**. Taken together, these data suggest that Hex deficiency did not affect the engraftment of donor-derived HCs, which was stable over time and ensured functional hemopoiesis.

**Figure 3.**
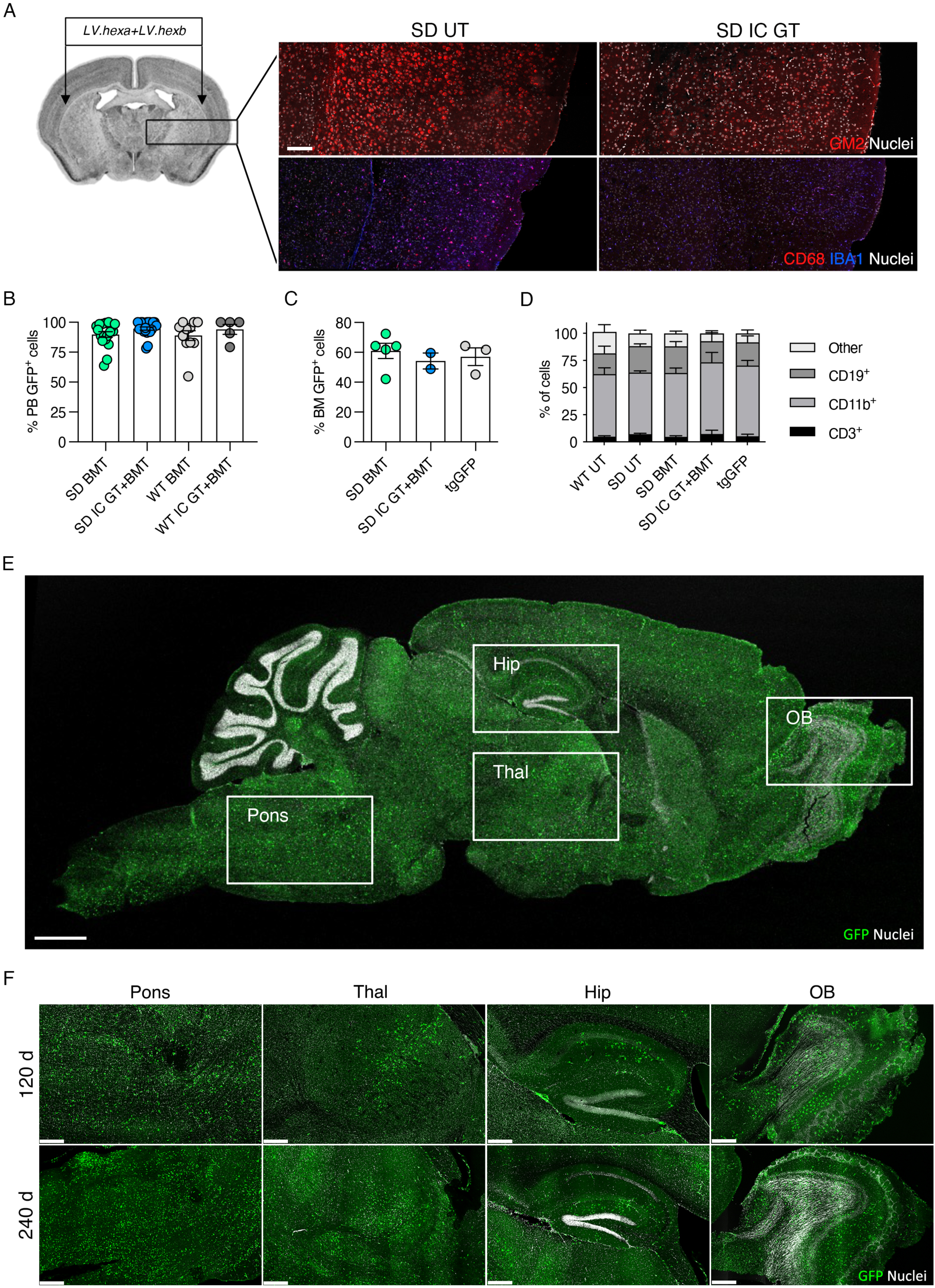
Transduction of neural cells upon IC GT and brain myeloid cell reconstitution upon BMT in SD mice. **(A)** Schematic of LV.hexa+LV.hexb injection and immunofluorescence (IF) confocal image showing clearance of GM2 storage and reduction in CD68+ in a region of SD brain tissue (box) close to the LV injection site. Upper panel: GM2, red; lower panel: CD68, red; IBA1, blue; nuclei are counterstained with Hoechst (white). Scale bar, 25 μm. **(B-C)** Percentages of donor-derived GFP+ cells in the peripheral blood (PB; B) and bone marrow (BM; C) of BMT- and IC GT+BMT-treated SD and WT mice evaluated at 30 d (PB) and 60 d (BM) after BMT. The percentage of GFP+ cells in the BM of tgGFP mice (donors) is shown for comparison. **(D)** Cellular composition of the BM analyzed 60 d after transplantation into BMT- and IC GT+BMT-treated SD mice and age-matched UT controls (WT, SD, and tgGFP mice). CD19, marker for B cells; CD11b, marker for monocytes, neutrophils, and NK cells; CD3, marker of mature T cells. Data represent the mean ± SEM; n= 3-13 mice/group. **(E)** IF image showing distribution of GFP+ cells in the whole brain of IC GT+BMT-treated SD mouse at 120 d. Boxes highlighting brain regions in (F) at higher magnification. **(F)** IF pictures showing GFP+ cells engrafted into olfactory bulb (OB), hippocampus (Hip), thalamus (Thal), and Pons/medulla (Pons) of IC GT+BMT-treated SD mice at 120 d and 240 d. In E and F: direct GFP fluorescence, green; nuclei counterstained with Hoechst, grey. Scale bars, 3.000 μm (E) and 500 μm (F).

Importantly, immunofluorescence (IF) analysis of brain tissues of BMT- and combination-treated mice at 120 d and 240 d showed the robust, widespread, and time-dependent engraftment of donor-derived GFP+ cells, which were distributed rostro-caudally within some areas of higher GFP+ cell density (i.e., olfactory bulbs, thalamus, pons, and medulla) (**Figure 3E-F**). A similar distribution of GFP+ cells was found in BMT- (analyzed at 120 d of age) and combination-treated SD mice (analyzed at 120 d and 240 d), suggesting that IC GT did not impact the modality of HC recruitment/engraftment in CNS tissues.

### Enhanced reconstitution of Hex activity in CNS tissues of combination-treated SD mice

To evaluate the efficacy of *in vivo* transduced neural cells upon IC GT and donor-derived HC myeloid progeny upon BMT and IC GT + BMT to the supply of functional enzyme in the CNS and periphery, we measured Hex activity in the telencephalon (TEL), cerebellum (CB), spinal cord (SC), cerebrospinal fluid (CSF), BM, liver, spleen, and sciatic nerves (SN) of treated and UT SD mice using two artificial substrates: 3 mM 4-methyl-umbelliferyl-N-acetyl-β-d-glucosaminide (MUG), used to determine total Hex activity (HexA, HexB, and HexS); and 3 mM 4-methyl-umbelliferone-6-sulfo-2-acetamido-2-deoxy-β-d-glucopyranoside (MUGS), used to determine HexA activity. We analyzed cohorts of treated animals at 120 d (IC GT, BMT, IC GT + BMT) and 240 d of age (IC GT + BMT) and included age-matched WT and Het groups as controls.

The CNS tissues of UT SD mice (120 d) displayed minimal residual enzymatic activity (≤3% and ≤10% of physiological levels assessed using MUG and MUGS, respectively) due the presence of the αα isoenzyme (**Figure 4A** and **Supplementary Figure 2A-D**). WT and Het mice showed a significant caudal-to-rostral increase in Hex activity in the CNS, with the highest values measured in the TEL compared to the CB and SC. Interestingly, we detected a significant age-associated increase in enzymatic activity in WT tissues (**Figure 4B; Supplementary Figure 2A-D**).

**Figure 4.**
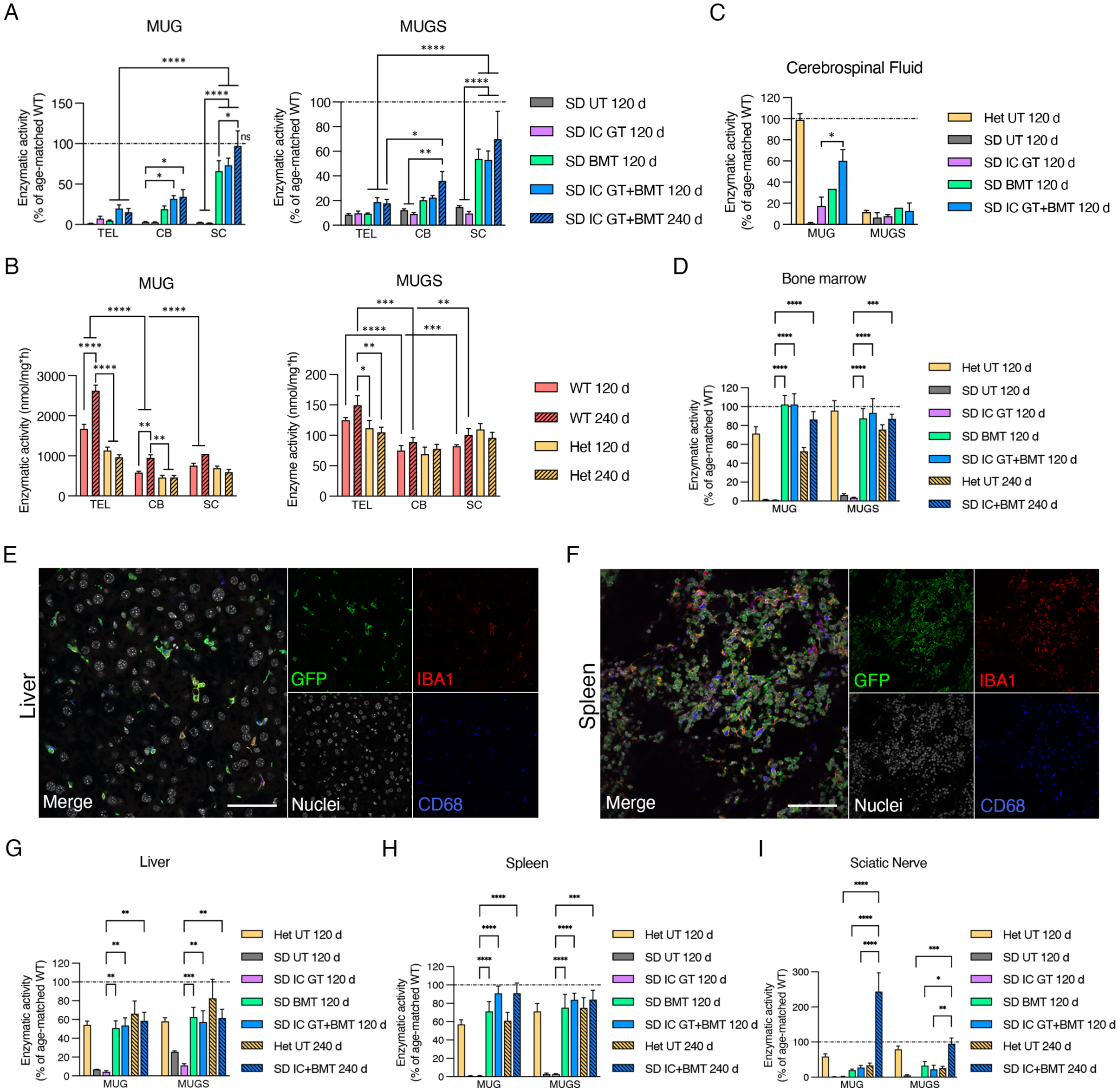
Hex enzymatic activity in CNS tissues, PNS, and periphery of treated SD mice and controls. **(A)** Enzymatic activity measured in the TEL, CB, and SC tissues of IC GT-, BMT-, and IC GT+BMT-treated SD mice and age-matched untreated (UT) SD mice at 120 d and 240 d. Enzymatic activity was measured as the degradation of the artificial substrates MUG (left graph) and MUGS (right graph) and expressed as percentage of age-matched UT WT control data. **(B)** Enzymatic activity (MUG, MUGS; expressed as nmol/mg/h) in the TEL, CB, and SC tissues of WT and Het mice at 120 d and 240 d. ^*^p < 0.01, ^**^p < 0.005, ^***^p < 0.001, ^****^p < 0.0001. **(C-D)** Enzymatic activity (MUG, MUGS; expressed as percentage of age-matched UT WT controls) measured in the cerebrospinal fluid (CSF; C) and bone marrow (BM; D) of treated SD mice (IC GT, BMT, and IC GT+BMT) and UT controls (Het, SD) at 120 d and 240 d. ^*^p < 0.05 (C); ^****^p < 0.0001, ^***^p < 0.001 SD IC GT 120 d vs all other treatments (D). Combined treatment vs WT UT, not significant. **(E-F)** Confocal IF images showing donor-derived GFP+ cells (green) expressing macrophagic marker Iba1 (red) and CD68 (blue) in liver (E) and spleen (F) of BMT-treated SD mice at 120 d. Scale bar, 50 μm. **(G-H-I)** Enzymatic activity measured in liver (G), spleen (H), and sciatic nerve (I) of treated SD mice (IC GT, BMT, and IC GT+BMT) and UT controls (Het, SD) at 120 d and 240 d. Enzymatic activity was measured as the degradation of MUG and MUGS and expressed as percentage of age-matched UT WT controls. ^**^p < 0.01, ^****^p < 0.0001 SD IC GT 120 d vs all other treatments (G-H). ^****^p < 0.0001, ^**^p < 0.01 SD IC GT+BMT 240d vs all other treatments (I). Combined treatment vs Het (liver) and WT (spleen), p>0.05. Data in A-D and G-I are expressed as the mean ± SEM; n = 3-9 mice/group. Data analyzed by 2-way ANOVA followed by Tukey’s multiple comparison test.

IC GT-treated SD mice analyzed at 120 d displayed a modest increase in enzymatic activity in the TEL (≈7% of WT-MUG) but not the CB and SC, suggesting the poor diffusion of Hex away from the site of injection. In the CNS of BMT-treated mice, enzymatic reconstitution was low in the TEL (≈5% of WT-MUG), increased in the CB (≈20% of WT/MUG), and significantly higher in the SC (≈65% of WT-MUG), likely reflecting caudal-to-rostral HC engraftment in the CNS tissues (**Figure 4A; Supplementary Figure 2A-C**). The similar enzymatic activity recorded in the TEL of IC GT- and BMT-treated SD mice highlighted the modest but stable enzymatic supply provided by neonatal IC GT in this region, in which HC-derived myeloid reconstitution was delayed compared to that in caudal regions. Combination-treated SD mice showed higher Hex reconstitution compared to BMT-treated mice in the TEL and CB (≈20% and ≈34% of WT activity-MUG, respectively) and a significant increase in the SC (≈73% of WT-MUG) (**Figure 4A; Supplementary Figure 2A-C**). The presence of robust Hex activity in the CSF of combination-treated SD mice (≈60% of WT-MUG) compared to IC GT- (≈18% of WT-MUG) and BMT-treated mice (≈34% of WT-MUG (**Figure 4A; Supplementary Figure 2A-C**) confirmed the production and release of functional enzyme by LV-transduced neural cells and donor HC-derived brain myeloid cells. The synergy of the treatments became even more evident in the combination-treated animals analyzed at 240 d of age, which showed stable Hex activity in the TEL and CB and physiological Hex levels in the SC (**Figure 4A; Supplementary Figure 2A-C**). Importantly, DEAE chromatography performed on TEL tissues showed normal patterns of HexA and HexB isoenzymes in the combination treated SD mice (**Supplementary Figure 2E**).

We next assessed the enzymatic reconstitution in the BM niche and the time-dependent contribution of donor HC-derived myeloid progeny in providing functional Hex to the PNS and periphery. The normalization of enzymatic activity measured at 120 d and 240 d in the BM of BMT- and combination-treated SD mice confirmed the high and stable peripheral chimerism and complete reconstitution of the hematopoietic system 2 months after the transplant (**Figure 4D; Supplementary Figure 2F**). The high numbers of GFP+CD68+ macrophages detected in the liver (**Figure 4E**) and spleen (**Figure 4F**) of BMT-treated SD mice at 120 d was in line with the observed rescue of Hex activity to heterozygous (liver) and WT enzymatic levels (spleen) (**Figure 4G-H; Supplementary Figure 2G-H**). Interestingly, Hex activity in the SN of BMT-transplanted mice was rescued to heterozygous (120 d), physiological, or supraphysiological levels (240 d) (**Figure 4I; Supplementary Figure 2I**), suggesting the kinetics of the PNS macrophage replacement was comparatively slow. Enzymatic activity in the peripheral tissues of IC GT-treated SD mice was similar to that of UT SD controls, concurrent with previous studies showing the CNS-specific targeting of this approach (32).

Overall, these results suggest there was complete repopulation of spleen and liver macrophages by donor HC-derived myeloid progeny 2 months after BMT in SD mice. In contrast, at least 4 months were needed for the significant repopulation of rostral brain regions, in which the requirement for enzymatic reconstitution was the highest. This timing is not very compatible with the rapid acceleration in disease progression experienced by these mice, unless a complementary source of early enzymatic supply (in this case provided by IC GT) is available. Interestingly, a similar timeframe was required for the full repopulation of the PNS by HC-derived macrophages and the maximum restoration of Hex activity.

### Correction of lysosomal expansion and clearance of GM2 storage in the CNS of combined-treated mice

We next investigated whether the enzymatic rescue provided by the treatments would lead to the amelioration/rescue of SD pathological hallmarks in CNS tissues. Increased expression of lysosomal-associated membrane protein 1 (LAMP1) is associated with lysosomal impairment in several LSDs (55). We discovered significant age- (120 d > 90 d > 30 d) and region-dependent (CB > SC > TEL) accumulation of LAMP1 protein in the CNS tissues of UT SD mice compared to the age-matched WT controls (4- to 10-fold higher than the WT levels; **Supplementary Figure 3**), which was reduced or normalized in the brain of treated SD mice at 120 d and 240 d (**Figure 5A**). These results suggest that the age-dependent reduction in LAMP1 expression in CNS tissues is related to the increased supply of functional Hex provided over time by the treatment.

**Figure 5.**
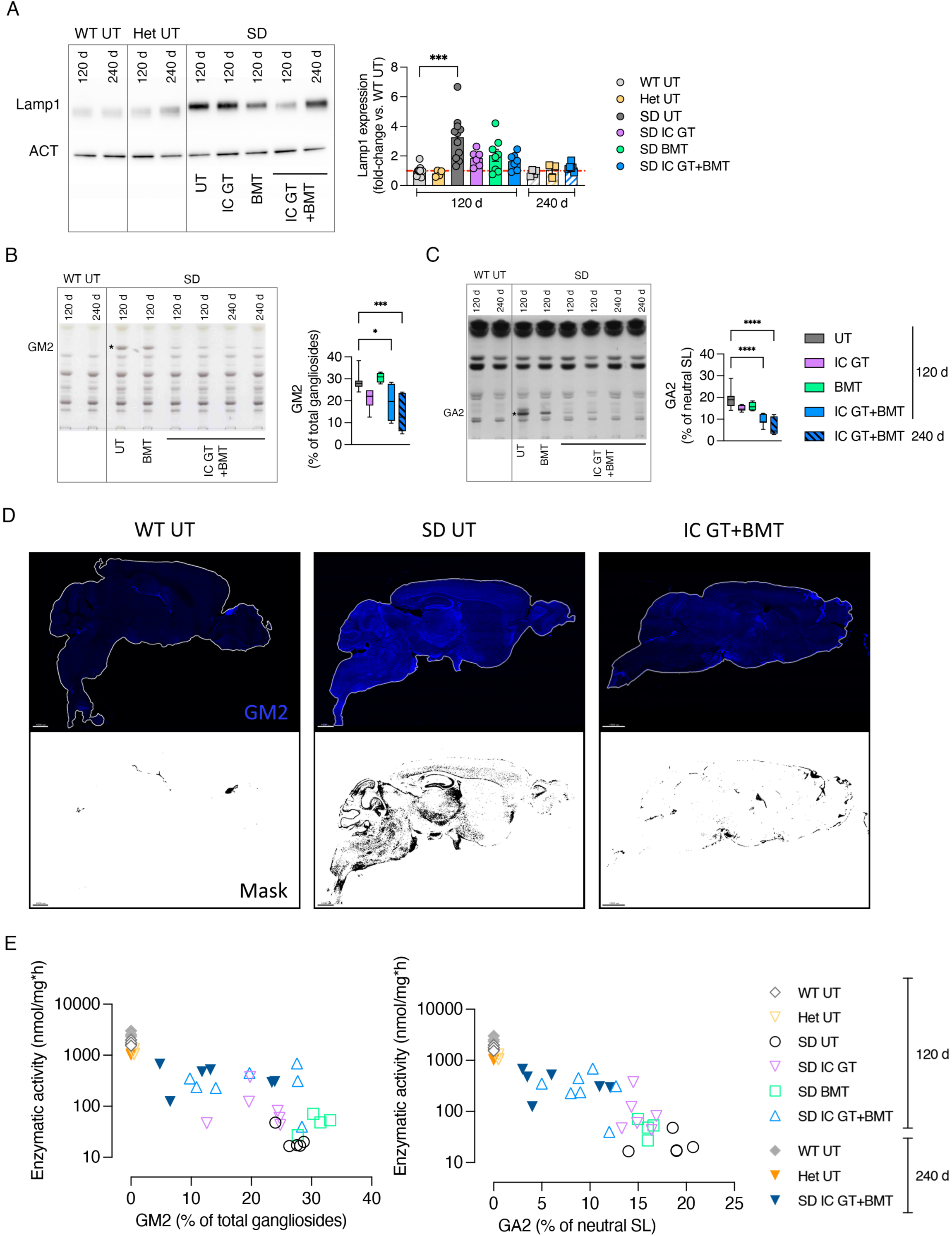
Rescue of LAMP1 expression and GM2 storage in treated SD mice. **(A)** Representative western blot analyses and relative quantification showing LAMP1 protein expression in whole brain lysate (TEL, CB) of treated mice (IC GT, BMT, and IC GT+BMT) at 120d and 240d and age-matched UT controls (WT, Het, and SD). Data are expressed as the mean ± SEM; n = 3-11 mice/group; one-way ANOVA followed by Kruskal–Wallis multiple comparison test, ^***^p < 0.001. **(B-C)** Representative HPTLC and relative quantification of aqueous phase (GM2, **B**) and organic phase (GA2, **C**) of total lipids obtained from whole brain lysate (TEL, CB) of BMT- and IC GT+BMT-treated SD mice (120 d and 240 d) and age-matched UT controls (WT, SD). Quantifications are expressed as percentage of total gangliosides (GM2) and percentage of total neutral sphingolipids (GA2). Data represent the mean ± SEM; n = 4-8 mice/group. One-way ANOVA followed by Dunnett’s multiple comparison test, ^*^p < 0.05, ^***^p < 0.001, ^****^p < 0.0001. **(D)** Representative IF pictures showing GM2 expression (anti-GM2 antibody; blue) and relative masks selection of immunopositive signal (black) in sagittal brain sections taken from IC GT+BMT-treated SD mice and UT controls (SD, WT) at 120 d. Magnification 10X; Scale bars, 1000 μm. **(E)** Correlation of Hex enzymatic activity (y-axis; MUG, nmol/mg/h) and GM2 or GA2 levels (x-axis; values expressed as percentage of total gangliosides or total neutral sphingolipids, respectively) in the TEL of treated SD mice at 120 d and 240 d and age-matched UT controls (SD, WT, Het). Each dot represents one animal. Treatment groups are shown in the legend.

Hex deficiency significantly increases the concentration of brain GM2 ganglioside and its asialo counterpart gangliotriaosylceramide (GA2), which are only present at trace concentrations in physiological conditions (56). We detected increased GM2 and GA2 contents in the brain tissues of UT SD mice at 30 d and 60 d of age, i.e., the asymptomatic stage, compared to WT controls. In 120-d-old fully symptomatic mice, the GM2 ganglioside and GA2 contents were 28.7 ± 4.2% and 19.5 ± 4.5% of the total gangliosides and total neutral sphingolipids, respectively (**Supplementary Figure 4A-B**). To evaluate the contribution of different therapies in preventing, delaying, and/or reducing the ganglioside store, we quantified the GM2 and GA2 in the brain tissue lysates of treated and UT SD mice compared with WT/Het controls. GM2 and GA2 storage was unchanged or slightly decreased in IC GT- and BMT-treated SD mice analyzed at 120 d of age. In contrast, the combination-treated mice showed GM2 and GA2 levels similar to those measured in asymptomatic 30-d-old UT SD mice. This reduction was even more apparent in mice analyzed at 240 d of age (**Figure 5B-C**). Qualitative IF analysis of the brain tissue of treated mice and controls with an anti-GM2 antibody confirmed the efficacy of the combined treatment in clearing ganglioside storage (**Figure 5D**), specifically in brain regions such as the olfactory bulbs, septum, and dentate gyrus, where GM2 accumulates first and most abundantly (**Supplementary Figure 4C**).

The extent of residual GM2 and GA2 stores in the brain tissues of treated mice correlated with the level of Hex activity (**Figure 5E**). The modest enzymatic supply provided by IC GT or BMT (<10% of WT-MUG) resulted in minimal or no reduction in storage levels. In contrast, 20-25% of the physiological Hex activity provided by combination-treatment resulted in a remarkable reduction in GM2 and GA2 levels, with a tendency for improvement in 240-d-old-treated mice, in which GM2 and GA2 levels were reduced to 40-50% of those detected in fully symptomatic UT SD animals (**Figure 5B-C-E**).

Taken together, these data conclusively showed that the enzymatic supply provided to CNS tissues by single-treatment IC GT or BMT fell below the threshold levels required for therapeutic benefit. The reduced, even if not normalized, storage levels in the combination-treated mice suggests that supplying low levels of functional enzyme to the neonatal brain can foster the long-term BMT-mediated therapeutic effects.

### Amelioration of neuroinflammation by combined treatment

Astrogliosis and microgliosis contribute to the onset and maintenance of the disease phenotype of several neurodegenerative diseases and may precede the full manifestation of extensive neurodegeneration and demyelination in SD (19, 57). The time-course qPCR and immunofluorescence (IF) analyses performed on CNS tissues of asymptomatic (30 d and 60 d) and fully symptomatic (120 d) UT SD mice and age-matched WT controls showed disease- and age-dependent upregulation of inflammatory cytokines (Mip-1a/chemokine ligand 3-CCL3, Rantes/CCL5) and macrophage (CD68) and astrocyte (glial fibrillary acidic protein; GFAP) markers (**Supplementary Figure 5A-B**). BMT and combined treatments, but not IC GT, rescued CCL3, CCL5, and CD68 expression (**Figure 6A**), suggesting the major anti-inflammatory contribution of HC-derived myeloid cells in the SD brain. The pronounced astrogliosis observed in UT SD mice was partially reduced by combined treatment, but not by BMT alone, at 120 d and was stabilized at 240 d, as assessed at the mRNA (**Figure 6A)** and protein level (**Figure 6B**).

**Figure 6.**
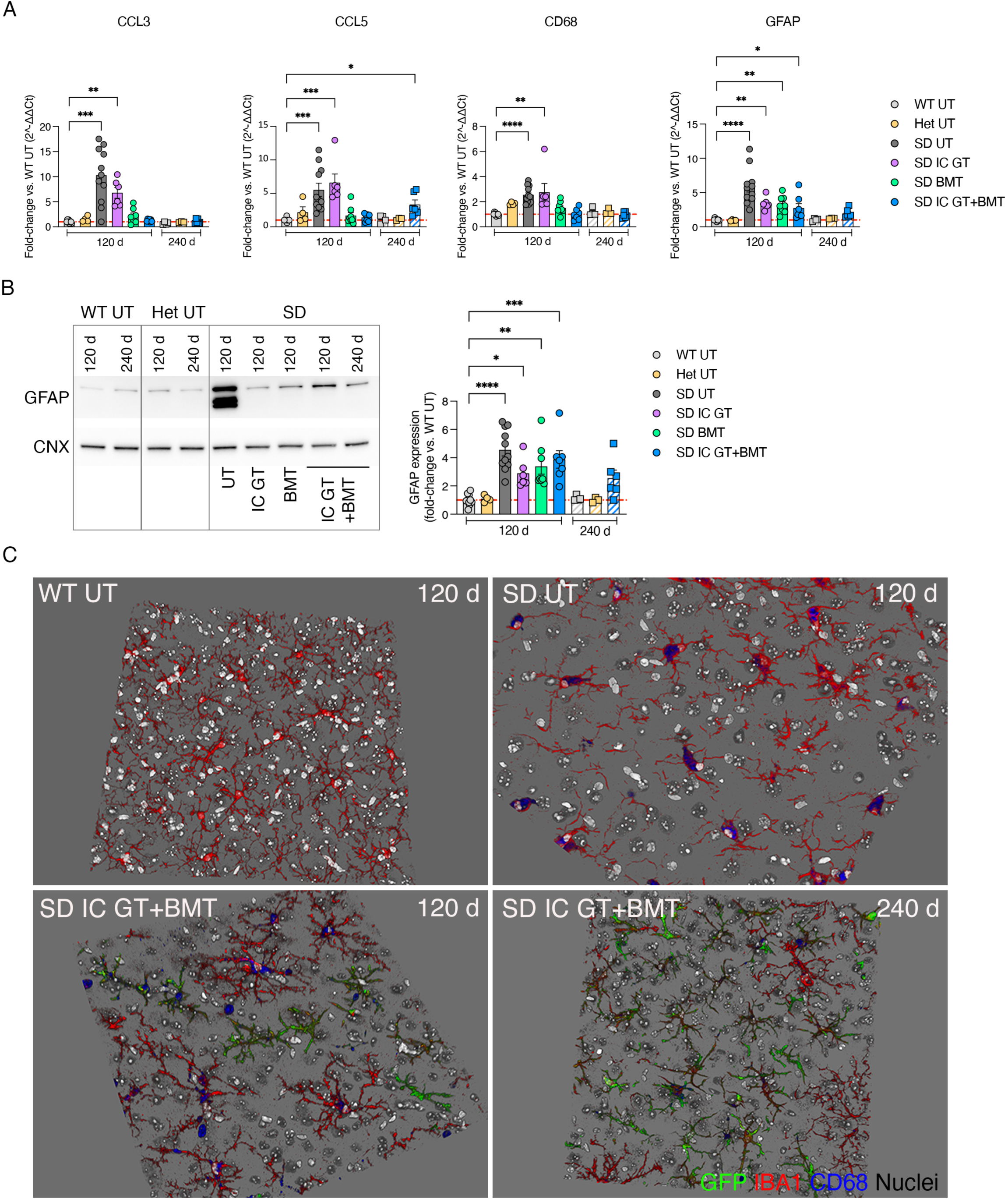
Reduction of neuroinflammatory markers in the CNS of treated SD mice. **(A)** Relative mRNA expression of neuroinflammatory cytokines (CCL3, CCL5, macrophage (CD68), and astrocytic (GFAP) markers in whole brain lysate (TEL, CB) of treated mice (IC GT, BMT, and IC GT+BMT) at 120 d and 240 d and age-matched untreated controls (WT, Het, and SD). Data are expressed as fold-change with respect to WT (set as 1) after normalization to *Gapdh* expression. **(B)** Representative western blot and relative quantification showing the expression of GFAP protein in treated mice (IC GT, BMT, and IC GT+BMT) at 120 d and 240 d and age-matched untreated controls (WT, Het, and SD). Data are expressed as fold-change to WT (set as 1) after normalization to calnexin (CNX) expression. Data in A and B represent the mean ± SEM; n = 3-11 animals/group. One-way ANOVA followed by Kruskal Wallis multiple comparison test, ^*^p < 0.05, ^**^p < 0.01, ^***^p < 0.001, ^****^p < 0.0001. **(C)** 3D projections of Z-stack images showing the presence of donor-derived GFP+ cells (green) and resident cells expressing Iba1 (microglia, red) and CD68 (macrophages, blue) in the TEL of IC GT+BMT-treated SD mice at 120 d and 240 d and UT controls (WT, SD). Nuclei counterstained with Hoechst, grey. Images were acquired at 40X magnification.

Qualitative IF analysis performed on brain tissues of treated SD mice and controls showed persistent reduction of CD68+ immunoreactivity in SD mice that received IC GT+BMT (120 d and 240 d; **Figure 6C**). Importantly, both resident and donor-derived (GFP+) Iba1+ cells in the combination-treated mice displayed a ramified, resting-like morphology, as opposed to the amoeboid shape that is associated with phagocytic/activated state observed in tissues from UT SD mice.

Overall, these results further highlighted the synergistic action of early IC GT and BMT against astrogliosis and microgliosis in the CNS of SD mice and in providing a long-term therapeutic advantage.

### Transient rescue of Purkinje cell degeneration by combined treatment

The evident ataxic phenotype of fully symptomatic SD mice prompted us to further investigate the potential cerebellar alterations. Qualitative IF analysis suggested there was less prominent GM2 storage in the CB of SD mice compared with other brain regions (see **Supplementary Figure 4C**), with an evident GM2+ signal in the granular and molecular layers and apparent modest storage in Purkinje cells (PCs) (**Figure 7A**). The altered morphology of PC dendritic trees and the altered localization of calbindin protein in the PCs of 30-d-old and 60-d-old asymptomatic SD mice compared to age-matched WT mice suggested the occurrence of neuronal dysfunction and/or ongoing neurodegeneration (**Figure 7B**). Loss of PC became evident in the 120-d-old symptomatic SD mice, which exhibited a ≈25% reduction in PC cells compared to the age-matched WT controls (**Figure 7C**). Importantly, the PC number, but not the calbindin mis-localization or of dendrite morphology, was transiently ameliorated in IC GT SD mice and normalized in BMT- and combination-treated animals (**Figure 7D**).

**Figure 7.**
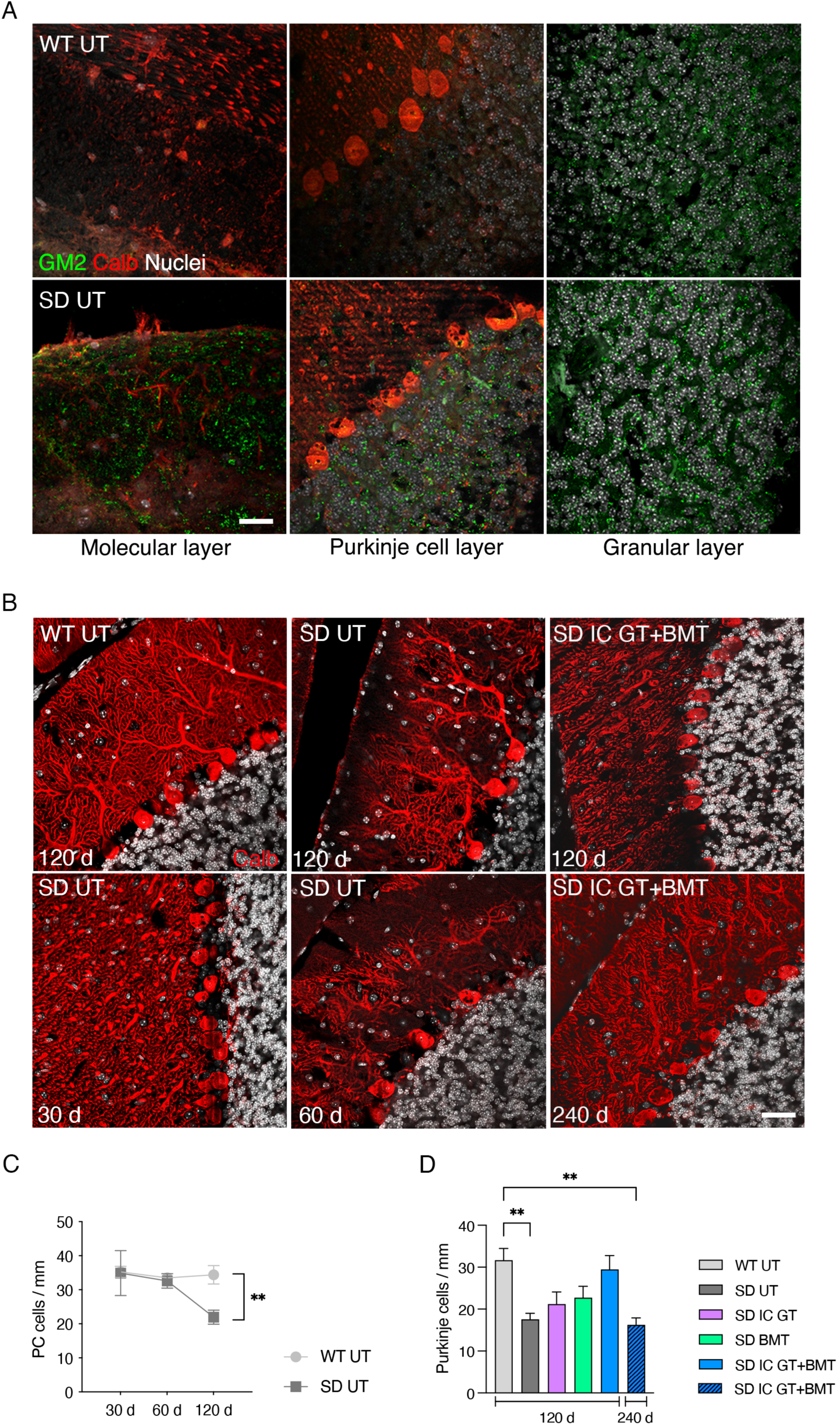
Effect of treatments on cerebellar pathology of SD mice. **(A)** Representative confocal pictures showing GM2 storage in the cerebellar molecular layer, Purkinje cell (PC) layer, and granular layer of UT WT and SD mice at 120 d. GM2, green; calbindin, red; nuclei are counterstained with Hoechst (white). Scale bar, 25 μm. **(B)** Representative confocal images showing calbindin+ PCs (red) in UT mice of different ages (WT:120 d; SD: 30 d, 60 d, and 120 d) and IC GT+BMT-treated SD mice at 120 d and 240 d. Calbindin, red. Nuclei counterstained with Hoechst, grey. Scale bar 50 μm. **(C)** Quantification of number of PCs in untreated (UT) SD and WT mice at 30 d, 60 d, and 120 d. Data represent number of PCs/mm and are expressed as the mean ± SEM; n = 1-10 mice/group. Two-way ANOVA followed by Bonferroni’s multiple comparisons test, ^**^p < 0.01. **(D)** Quantification of the number of PCs in treated mice (IC GT, BMT, and IC GT+BMT) at 120 d and 240 d and age-matched UT controls (WT, SD). Data represent number of PCs/mm within all cerebellar lobules of each brain section (3-4 sagittal sections per mouse). Data are expressed as mean ± SEM; n= 4-6 mice/group; one-way ANOVA followed by Kruskal Wallis multiple comparison test, ^**^p < 0.01.

### Partial rescue of myelin defects by combined treatment

In addition to the SD effects on neurons, defects in myelin structure and composition have been described in the CNS of SD animal models (56, 58–60) and patients (61). To assess whether single and combined treatments are able to preserve and/or rescue myelin integrity in CNS tissues, we performed morphometric analysis of SC and optic nerves. We did not observe any defects in the myelin or axons of SD mutant SN at 30 d or 120 d, confirming previous findings in the PNS (62, 63).

Semi-thin sections and ultrastructural analyses of cervical and lumbar SC regions of UT SD mice at 120 d revealed the presence of several fibers displaying demyelination and myelin degeneration. The total number of myelinated fibers and the axonal diameter distribution were similar in the mutant and controls (data not shown). However, the percentage of demyelinated/degenerated fibers within the total myelinated fibers was unchanged after BMT or IC GT+BMT treatment (**Figure 8A-B**). Of note, the similar percentage of demyelinated/degenerated fibers observed in the combination-treated SD mice analyzed at 120 d and 240 d suggests that this strategy might inhibit disease progression and prevent further myelin deterioration (**Figure 8A-B**).

**Figure 8.**
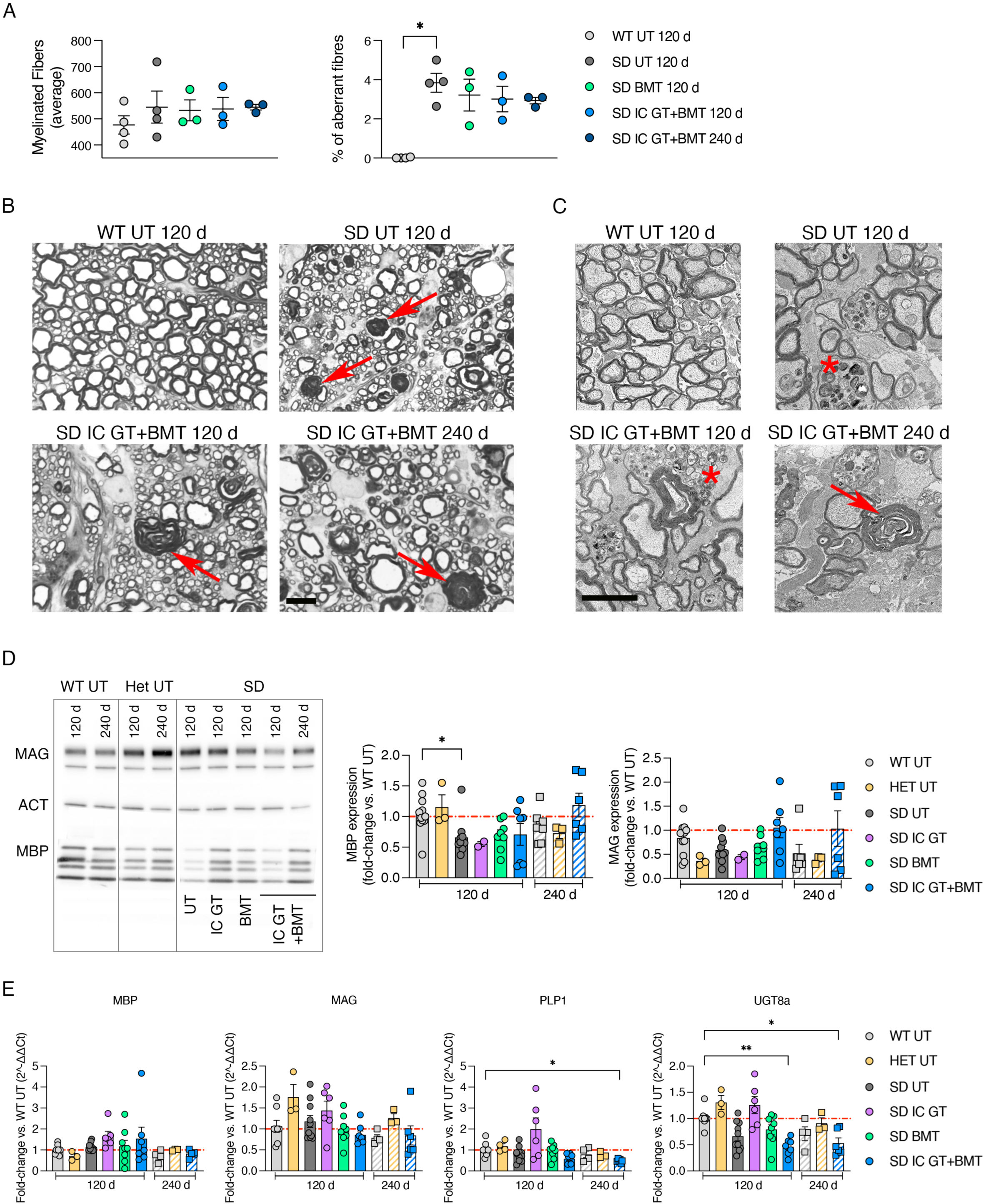
Analyses of myelin compartment in treated SD mice. **(A)** Quantification of myelinated fibers and percentage demyelinated/degenerated (aberrant) fibers of total fibers in spinal cord sections from treated mice (IC GT, BMT, and IC GT+BMT) at 120 d and 240 d and age-matched UT controls (WT, SD). Data are expressed as mean ± SEM; n = 3-4 mice/group; nonparametric one-way ANOVA followed by Dunn’s multiple comparison, ^*^p < 0.05. **(B)** Semithin section analysis of lumbar SC of IC GT+BMT-treated SD mice at 120 d and 240 d and UT controls (WT, SD). Red arrows indicate aberrant myelin. Scale bar, 10 µm. **(C)** Electron microscopic images of optic nerves in treated SD and UT controls at 120 d and 240 d. Red arrows indicate aberrant myelin and asterisks mark macrophages. Scale bar, 2 µm. **(D)** Representative western blot analyses and relative quantification showing MBP and MAG protein expression in whole brain lysate (TEL, CB) of treated mice (IC GT, BMT, and IC GT+BMT) at 120 d and 240 d and age-matched UT controls (WT, Het, and SD). Data are expressed as fold-change to WT (set as 1) after normalization to -actin expression. Data represent the mean ± SEM; n = 3-10 mice/group. One-way ANOVA followed by Kruskal–Wallis multiple comparison test, *p < 0.05. **(E)** Expression of myelin-related genes (*Mbp, Mag, Plp1*, and *Ugt8a*) in whole-brain lysate (TEL, CB) of treated mice (IC GT, BMT, and IC GT+BMT) at 120 d and 240 d and age-matched UT controls (WT, Het, and SD). Data are expressed as fold-change with respect to the WT (set as 1) after normalization to *Gapdh*. Data represent mean ± SEM; n = 2-12 mice/group. One-way ANOVA followed by Kruskal-Wallis multiple comparison test, ^*^p < 0.05, ^**^p < 0.01.

Electron microscopic analyses of the optic nerves of UT SD mice (120 d) revealed macrophage infiltration and signs of axonal degeneration (organelle swelling and accumulation of storage materials). Contrary to what was observed in the SC sections, this phenotype was not ameliorated by treatment but was aggravated in the combination-treated SD mice at 240 d, at a time when the loss of myelinated fibers was observed (**Figure 8C**).

To support the morphological observations, we first analyzed the expression of myelin proteins in brain lysates from UT and treated SD mice at 120 d (IC GT, BMT, IC GT+BMT) and 240 d (IC GT+BMT) on western blot, including the age-matched WT and Het mice controls (**Figure 8D**). Quantification of the immunoblot bands showed a 40-50% reduction in myelin basic protein (MBP) and myelin-associated glycoprotein (MAG) expression in UT SD mice as compared to WT controls at 120 d of age. Both BMT and combined treatments (but not IC GT) rescued MBP expression, which was stable in the combination-treated SD mice at 240 d. We observed a 50% reduction in MAG expression in 240-d-old compared to 120-d-old WT mice. The combination-treated SD mice analyzed at 240 d showed similar MAG levels to the age-matched WT controls. In line with previous reports (60), we detected similar expression levels of the myelin genes *Mbp, Mag, Plp1* (ProteoLipid Protein 1), and *Ugt8a* (UDP galactosyltransferase 8A) in the brain tissues of UT SD mice and WT controls, and these levels were not significantly affected by the treatments (**Figure 8E**).

Overall, these data suggest that a disease-associated alteration in brain myelin protein composition can be rescued by combined therapy in SD mice, whereas it remains relatively refractory to correction by isolated therapies.

## DISCUSSION

In this work, we showed the complementary mode of actions and synergistic effects of LV-mediated intracerebral GT and BMT in providing a significant therapeutic enzymatic supply and counteracting multiple pathological hallmarks in the brain tissues of SD mice, which recapitulated the severity and rapid progression of early-onset forms of SD.

Combination-treated mice preserved their normal body weight and motor functions until 150 days of age, a time at which all untreated/IC GT-treated SD mice and the majority of BMT-treated SD mice were deceased. Importantly, combination-treated SD mice analyzed at 240 d (2-fold more than the average survival and chosen as the experimental endpoint) showed only minor disease manifestations, suggestive of the prolonged efficacy of the combined treatment. The increase in survival reported here for the combination-treated mice was similar or lower to that reported by previous studies in which the authors co-delivered two AAVs expressing the and subunits into the brain parenchyma (23, 60, 64, 65) or systemically injected bicistronic AAV9 constructs (66, 67). Despite the promising pre-clinical data sustained the development of AAV GT, there are still several drawbacks associated with both intracerebral and systemic AAV delivery, including a failure to correct early pathological hallmarks (e.g., myelin defects) (22, 56), the neurotoxicity associated with transgene overexpression in brain cells (27), and the potential liver and heart toxicity associated with off-target transgene expression (66–69). These considerations, together with the known immunogenicity of AAVs (70), dictate the need to accurately monitor patients enrolled in the first in-human AAV GT trials for GM2 gangliosidosis (NCT04798235, NCT04669535) and provide the rationale for exploring alternative/complementary therapeutic strategies.

Therapeutic improvements in GM2 gangliosidosis rely on the rescue of Hex enzymatic activity to clear or counteract ganglioside storage in affected tissues, specifically in the CNS. The comparative Hex activity among different brain areas under physiological conditions has not been comprehensively investigated. The 2.5-fold higher Hex activity measured in the TEL compared to that in the CB and SC tissues of adult WT mice suggests that the enzymatic requirement is higher in the rostral versus the caudal CNS regions. The minor increase in Hex activity (7% of the normal value) in the TEL of IC GT mice and the modest amount of secretion/transport is peculiar to Hex, as other lysosomal enzymes (i.e., ARSA and GALC) are found throughout CNS tissue upon local intraparenchymal LV injection and supply 30-100% of normal enzymatic activity (31, 32). Despite the modest increase, the enzyme provided by IC GT is available from a few days post-neonatal injection (71) and is stable over the months before the BMT, and within the timeframe required for full BM-derived myeloid cells engraftment in the CNS. Thus, it likely plays a key role in limiting GM2 storage build up and delaying pathological progression. Upon BMT, donor BM-derived myeloid cells engraft into the CNS, providing additional enzyme and trophic support and reconstituting the myeloid cell population in the PNS and peripheral organs, which are not targeted by intraparenchymal LV Hex injection. The use of full myeloablation conditioning with busulfan (54) accounted for the robust and stable hematopoietic chimerism achieved in SD mice upon BMT in this study (90-100%) compared to those in previous studies in which the engraftment was low or not assessed (18, 19, 50, 72). The analysis performed at 120 d showed complete reconstitution of enzymatic activity in the BM of transplanted SD mice— indicative of the complete repopulation of the hematopoietic compartment—and the significant and stable rescue of enzymatic activity in the liver and spleen (40-80% of WT), indicating the presence of functional donor-derived tissue macrophages behaving as a long-term source of Hex enzyme. Interestingly, we observed slower kinetics for the Hex activity rescue in the SN, suggesting that at least 6 months are necessary to achieve full repopulation of the PNS and peripheral organs by donor-derived cells. This slow repopulation may, in part, explain the limited efficacy of the allogeneic hematopoietic stem cell transplantation (HSCT) in rescuing the severe and progressive peripheral neuropathy associated with rapidly progressive forms of neuronopathic LSDs (73, 74).

In line with previous studies (51, 72), we found a time-dependent caudal-to-rostral distribution of donor-derived myeloid cells in the CNS tissues of BMT- and combination-treated mice. Accordingly, enzymatic reconstitution was achieved earlier in transplanted SD mice and at higher levels in the SC, with the TEL showing the lowest Hex activity with respect to physiological levels. The increase in enzymatic activity measured in combination-treated animals at 240 d of age further confirmed that the infiltration of donor-derived myeloid cells into the CNS is a slow and progressive process, and it supported the rationale for using combined approaches to delay or halt the progression of the early-onset forms of GM2 gangliosidosis. Indeed, the progressive rescue of enzymatic activity provided by combined therapy, but not IC GT or BMT in isolation, normalized LAMP1 expression and strongly reduced GM2 and GA2 storage to levels comparable to those found in 30-d-old asymptomatic UT SD mice. Importantly, our analysis showed a clear correlation between enzymatic activity and residual ganglioside storage in the TEL, indicating that at least 20-25% of normal enzymatic activity (measured from the total tissue lysate and CSF) is needed to ensure the 50-70% reduction in GM2 and GA2 storage that distinguishes the therapeutic effects of combined treatments from those of single treatments. Our results concur with previous studies showing 30-60% of normal Hex enzymatic activity is required for a 80-90% reduction in GM2 storage in the CNS tissues of affected mice (67)(75), and with reasonable accuracy, they indicate on the threshold of enzymatic activity that, if measured in the CSF of patients undergoing experimental treatments, could be associated with therapeutic benefit.

The complementary kinetics of the enzyme provided by IC GT and BMT, as well as the modality of action of these treatments, largely account for their synergy when combined. Reactive astrocytes, resident macrophages/microglia, and blood-borne immune cells are critical cellular components in mediating neuroinflammation (76–78), which is a hallmark of different neurodegenerative disorders and has been linked to several LSDs, including GM2 gangliosidosis (79). Our results indicate there is an important inflammatory response in the regions of SD mouse brains where GM2 storage is more prominent. The local source of enzyme provided by IC GT is insufficient to counteract progressive neuroinflammation. In contrast, animals transplanted with BMT showed reduced expression levels of inflammatory-related genes, confirming the immunomodulatory and anti-inflammatory properties of BM-derived cells engrafted into the CNS (18, 80). Long-lived combination-treated animals showed normalized inflammatory conditions and reduced astrogliosis, confirming the synergistic effect and long-lasting efficacy of the sequential treatments.

While our results support the efficacy of combined treatments in providing a timely enzymatic supply and the rescue of disease-associated hallmarks in multiple affected tissues and organs, they also highlighted some aspects of the complex SD pathology that are refractory to correction. Neuronal apoptosis and degeneration have been described in specific regions of GM2-gangliosidos animal models and patients (23, 81). However, the molecular mechanisms whereby GM2 accumulation in neurons triggers neurodegeneration remain unclear. Cerebellar pathology has been described in juvenile and late onset TSD and SD patients (82–85), in a sheep model of GM2 gangliosidosis (86), and in a mouse model of GM2 activator deficiency (87), but it has been poorly investigated in SD mice. We observed altered cerebellar PC morphology in symptomatic SD mice, which are clearly ataxic. Interestingly, moderate hindlimb ataxia and weakness persists in long-lived combination-treated SD mice. Further investigations are warranted to understand the cause-and-effect relationships of GM2 storage and calbindin expression and localization in PCs and other cerebellar cell types involved in the neural circuits that regulate motor coordination. Combined treatments and BMT transiently counteracted PC loss without completely rescuing cell morphology and normalizing calbindin expression. The direct cerebellar injection of therapeutic vectors could be more effective in correcting these pathological hallmarks.

Defects in myelin structure and composition have been described in the CNS of GM2 gangliosidosis animal models (mice, cats, dogs, and sheep) (56, 58–60, 88, 89) and patients (61). However, it is still unclear whether the myelin defects are a consequence of neurodegeneration or whether they have a primary role in the pathogenesis, as suggested by their early occurrence in SD mice (60). Here, we have shown that the progressive hypo/demyelination and infiltration of inflammatory cells into the SC tissues of 120-d-old SD mice is stabilized, if not normalized, by combined IC GT and BMT treatments. These effects were not observed in the optic nerve, where extensive fiber loss was present at 240 d post treatment. Further analyses are needed to investigate the role of region-specific demyelination in disease progression, to establish whether it is a consequence of or a contributing factor to the pathological phenotype, and to discover effective strategies to counteract the phenomenon.

In conclusion, the results of this study provide proof-of-concept evidence for the feasibility and efficacy of combining LV-mediated IC GT and BMT to target the CNS, PNS, and peripheral tissues with appropriate timing to counteract rapid disease progression in a relevant murine model of GM2 gangliosidosis. Our study highlighted that the short-term advantage provided by early (even if only modest) enzymatic supply in CNS tissues through IC GT enhances the long-term benefits of HC-derived myeloid progeny, which provides a reservoir of functional enzyme in all affected tissues and counteracts the detrimental neuroinflammation that current *in vivo* IC GT approaches fail to fully correct. We also identified the threshold for tissue/CSF enzymatic activity that is required to achieve a reduction in brain GM2 storage, which correlates with improved benefit, with important implications for the interpretation of current and future experimental trials. In terms of clinical translation, the use of autologous hematopoietic stem cells genetically modified by LVs to express Hex enzymes (36) may overcome the current limitation of allogeneic BMT or HSCT, overall improving the safety and increasing the benefits of the proposed combined strategy. Of note, *ex vivo* HSC GT has recently received full EU marketing authorization for the treatment of early-onset metachromatic leukodystrophy patients (using autologous CD34+ cells transduced with an LV encoding the ARSA gene - Libmeldy) (90, 91) and is in advanced stages of clinical testing for other neurodegenerative LSDs (92)(74). The optimization of combined LV-mediated *in vivo* and *ex vivo* GT protocols could address the unmet medical needs of patients with GM2 gangliosidoses, as well as, potentially, other LSDs for which therapeutic options are absent or insufficient.

## MATERIALS AND METHODS

### Animals

SD mice (hexb-/-) were generated as previously described (17). Mice were bred via heterozygosis, and Hexb+/+ and hexb+/- littermates were used as controls. CAG GFP transgenic mice [C57BL/6-Tg (CAG-EGFP)1Osb/J] were purchased from Jackson Labs (Bar Harbor, Maine, 04609, USA). CAG GFP mice have an enhanced GFP (EGFP) cDNA under the control of a chicken β-actin promoter and cytomegalovirus enhancer and show widespread EGFP fluorescence. Mouse colonies were maintained in the animal facility at the San Raffaele Scientific Institute, Milano, Italy.

### Lentiviral vectors

We used monocistronic third-generation LVs expressing cDNA encoding murine hexa or murine hexb (LV.mA, LV.mB) or GFP cDNA (LV.GFP) under the control of the human PGK promoter. LVs were cloned, produced, and titrated as previously described (53):

- LV.mA. Titer: 0.68 × 10^9^ TU/ml, infectivity: 3.78 × 10^4^ TU/ng
- LV.mB. Titer: 1.15 × 10^9^ TU/ml, infectivity: 7.47 × 10^4^ TU/ng
- LV.GFP. Titer: 1.1 × 10^10^ TU/ml, infectivity: 1.5 × 10^5^ TU/ng

### Treatments

#### Intracerebral gene therapy (IC GT)

SD mice and controls were injected with LV preparations at 2 d. Pups were anaesthetized in crushed ice and placed on a refrigerated stage. LVs (4 × 10^6^ TU/μl) were injected bilaterally into the external capsule (EC) under stereotactic guidance using a Hamilton syringe (gauge 33; Model 1701 RN Neuros Syringe, #65460-05 10µL; Hamilton, Reno, Nevada, USA). Stereotactic coordinates (mm from the frontal cerebral vein): AP +1, ML +2, DV −1. After injection, the pups were placed on a heating pad, monitored for 15 min, and then replaced in their cage with the mother.

#### Bone marrow transplant

Total BM was flushed from the tibias, femurs, and humeri of 4- to 8-week-old tgGFP mice as previously described (51). One tgGFP mouse served as the donor for six recipient mice. Cells were suspended in PBS (5 × 10^6^ cells/200 μl) and immediately injected into the tail vein of 60-day-old busulfan-conditioned SD mice (125 mg/kg for 4 days) (93). Survival after the procedure was 100%. Part of the BM suspension was collected for enzymatic and cytofluorimetric activity, and 100% of BM cells isolated from the tgGFP mice were GFP+ and expressed physiological levels of β-hex activity.

#### Combined treatment

60-day-old mice that had been treated with IC GT on PND2 received BMT the day after completion of the myeloablation protocol by i.p. busulfan injection (125 mg/kg for 4 d). Mice dying within 2-3 weeks after busulfan conditioning were excluded from subsequent analyses. Animals were euthanized at 120 days of age (average lifespan of untreated SD mice), 240 days of age (chosen as endpoint of the study), or when they reached the human disease endpoint (weight loss >80% with respect to age-matched WT mice or inability to eat and drink). Untreated SD, Het and WT littermates were included as controls.

### Tissues collection and processing

Mice were euthanized after anesthetics overdose (ketamine/xylazine) via intracardiac perfusion of the descending aorta with 0.9% NaCl. Brain hemispheres were separated, and one hemisphere was cut to separate the TEL from the CB; the tissues were immediately frozen and used to measure enzymatic activity (94). The other hemisphere was cut into two equal sagittal slices: one was post-fixed in 4% paraformaldehyde (PFA) in PBS and used for IF, the other was smashed through a cell strainer in PBS, washed, pelleted, and quickly frozen for later biochemical and molecular assays (thin-layer chromatography, and RNA and protein extraction). Whole SC was collected and sagittally halved, and the SN, spleen, and liver were isolated and split in half. For each of these tissues, one half was quickly frozen in liquid nitrogen for enzymatic activity analysis, and the other half was post-fixed in 4% PFA in PBS for IF analysis. Total BM was flushed from tibias, femurs and humeri and collected for enzymatic activity and cytofluorimetric analysis. We collected the CSF from the cisterna magna immediately prior to euthanasia using a glass capillary (only in mice at 120 d).

### Immunofluorescence

Free-floating vibratome sections (40-µm thick) were incubated with blocking solution (10% normal goat serum [NGS] + 0.3% Triton X-100 in PBS) for 1 h at RT and then incubated overnight at 4°C with primary antibody diluted in blocking solution. After a thorough three washes of 5 min each, species-specific fluorophore-conjugated secondary antibodies diluted in 10% NGS in PBS were added. Coverslips and tissue sections were counterstained with 4’, 6-diamidino-2-phenylindole (Hoechst; Roche, Rotkreuz, Switzerland) for nuclei, washed in 1x PBS, collected, and mounted on glass slides using Fluorsave (CALBIOCHEM; San Diego, CA, USA).

The following primary antibodies were employed: chicken anti-GFP (1:500; ab-13970, Abcam, Cambridge, UK); glial fibrillary acidic protein (GFAP; polyclonal 1:1000, ZO334, DAKO-Agilent, Santa Clara, CA, USA; monoclonal 1:1000, MAB 3402, Millipore, Burlington, Massachusetts, United States); rabbit anti Iba1 (1:100; 019-19741, Wako Chemicals, Neuss, Germany); rat anti CD68 (1:200; clone FA-11/MCA1957, Bio-Rad Laboratories Inc, Hercules, CA, USA); rabbit anti LAMP1 (1:200, AB24170; Abcam, Cambridge, UK); rat anti LAMP1 hybridoma (1:300; 1D4B, DSHB at The University of Iowa, USA); mouse anti GM2 (1:500 A2576; TCI EUROPE); rabbit anti Calbindin (1:1000, CB-38A Swant, Burgdorf, Switzerland).

The secondary antibodies were from Thermo Fisher Scientific (Waltham, MA, USA): ALEXA 488 (1:1000; anti-rabbit A11008, anti-mouse A11001, anti-chicken A11039); ALEXA 546 (1:2000; anti-mouse A11039, anti-rabbit A11010); ALEXA 633 (1:500; anti-rat A21094, anti-mouse A21050).

### Image acquisition

Confocal images were acquired at different magnifications with a Leica TCS SP8 confocal microscope (Leica, Wetzlar, Germany) or a MAVIG RS-G4 upright confocal microscope (MAVIG GmbH Research, Munich, Germany). Data were analyzed with Fiji software (ImageJ, U. S. National Institutes of Health, Bethesda, Maryland, USA) (95), LasX software (Leica Application Suite X; RRID:SCR_013673), and Imaris viewer software (Oxford Instruments, Abingdon-on-Thames, UK). Images were imported into Adobe Photoshop 2021 for brightness and contrast adjustments and to merge the channels.

### Quantification of Purkinje cell number

Confocal images were acquired at 20X magnification using a MAVIG RS-G4 upright confocal microscope. We counted the number of PCs within the lobules of each section (3-4 sagittal sections per mouse) and related this value to the length of the relevant PC layer, as measured in micrometers using Fiji software (95).

### Cytofluorimetric analyses of peripheral blood and bone marrow samples

Peripheral blood (PB) was collected from BMT-treated mice and untreated controls, and 20 μL of each sample was incubated for 30 min at 4°C with rat anti-mouse CD45-VioBlue (Miltenyi Biotec, Bergisch Gladbach, Germany; Cat. 130-110-802), then for 15 min with ammonium-chloride-potassium (ACK) buffer for RBC lyses. Samples were centrifuged and suspended in FACS buffer (PBS + 5% FBS + 1% BSA). Then, the cells were analyzed for CD45 and GFP positivity using a flow cytometer (Canto II, BD Biosciences, Franklin Lakes, New Jersey, USA), and the data were analyzed using the FlowJo software (Ashland, Oregon, USA). The remaining aliquots of the PB samples aliquots were analyzed using an automated hemocytometer (IDEXX ProCyte Dx Haematology Analyzer, IDEXX Laboratories, Westbrook, ME, USA) to determine the complete blood count variables.

BM samples from donor and transplanted mice were resuspended in FACS buffer for 15 min and stained with CD45-BV510 (BD Biosciences, Franklin Lakes, New Jersey, USA; Cat. 563891), CD11b-APC (BD Biosciences; Cat. 553312), CD19-PE (BioLegend, San Diego, CA, USA; Cat. 152407), and CD3-FITC (BioLegend Cat. 100203) for 20 min; the GFP signal was measured by direct fluorescence. After washing in PBS, pellets were resuspended in 300 μl of PBS. Cells were analyzed by flow-cytometry using a CANTO instrument (Canto II, BD Biosciences). All data were analyzed using the FlowJo software.

### Morphological analysis of spinal cord and optic nerve

Semithin section analysis of SCs was performed as reported previously (96, 97). To perform morphometric analysis, digitalized images of SC cross-sections were obtained from similar regions (cervical and lumbar) with a 100X objective and Leica DFC300F digital camera. At least 10 images per animal were analyzed with Leica QWin software (Leica Microsystem, Wetzlar, Germany), and the total number of fibers, number of fibers displaying myelin degeneration, and the axon diameter distribution were calculated. Ultrastructural analysis was performed on optic nerves with 20-40 images per animal, which were acquired using the TALOS L120C transmission electron microscope (Thermo Fisher Scientific).

### Determination of Hex activity and isoenzyme composition

The β -Hex activity and chromatographic profiles were measured as previously described (53). Briefly, hexosaminidase activity was determined using MUG and MUGS substrates dissolved in 0.1 M citrate/0.2 M disodium phosphate buffer at pH 4.5. The enzymatic reactions were performed using 50 μl of test sample incubated with 100 μl of substrate at 37°C. All reactions were stopped by adding 2.850 ml of 0.2 M glycine/NaOH, pH 10.6. The fluorescence of the liberated 4-methylumbelliferone was measured on a Perkin Elmer LS50B spectrofluorometer (Waltham, Massachusetts; λ excitation 360 nm, λ emission 446 nm).

### Gene expression analyses

Total RNA from CNS tissues was extracted using RNeasy Lipid Tissue with Qiazol (Qiagen, Hilden, Germany) in accordance with the manufacturer’s protocol. The quantity of RNA was determined using a 260/280 nm optical density (OD) reading on a NanoDrop ND-1000 Spectrophotometer (NanoDrop, Pero, Italy). mRNA reverse transcription (RT) was performed using the QuantiTect reverse transcription kit (Qiagen) in accordance with the manufacturer’s protocol. Quantitative (q)PCR was performed in Optical 96-well Fast Thermal Cycling Plates on Viia7 (Thermo Fisher Scientific) using the following thermal cycling conditions, 1 cycle at 95°C for 5 min, 40 cycles at 95°C for 15 s and 60°C for 1 m, using Universal PCR Master Mix and TaqMan Gene Expression Assays (Thermo Fisher Scientific). SDS 2.2.1 software was used to extract raw data. The relative expression of mRNA for the target genes was calculated using the 2−ΔΔCt method.

Commercial probes and primers (ThermoFisher Scientific): *Gapdh* (glyceraldehyde 3-phosphate dehydrogenase), Mm99999915_g1; *Cd68*, Mm03047343_m1; *Ccl3*, Mm00441259_g1; *Ccl5*, Mm01302427_m1; *Gfap*, Mm01253033_m1; *Mbp*, Mm01266402_m1; *Mag*, Mm00487538_m1; *Plp1*, Mm00456892_m1; *Ugt8a*, Mm00495930_m1.

### Western blot analyses

Protein content was determined through a DC protein assay (Bio-Rad) following the manufacturer’s protocol. After boiling for 5 min in sample buffer, samples containing 5-20 µg of protein were separated on 4-12% acrylamide gel SDS-PAGE electrophoresis before antigen detection with the primary and secondary antibodies listed below. Immunodetection was performed using the Amersham ECL Plus kit (GE Healthcare, Little Chalfont, Buckinghamshire, UK). Primary antibodies: rabbit anti-LAMP1 (1:100; AB24170, Abcam); rabbit anti GFAP (1:100.000; ZO334, DAKO); mouse anti-MBP (1:4.000; clone SMI 94/SMI 99, BioLegend); rabbit anti-MAG (1:1.000; 34-6200, Thermo Fischer Scientific); rabbit anti-calnexin (1:5.000; C4731, Sigma-Aldrich, St. Louis, MO, USA); mouse anti-actin (1:10.000; sc-1616r, Santa Cruz Biotechnology, Dallas, TX, USA). Secondary antibodies (Chemicon, Temecula, CA, USA): goat anti rabbit HRP (1:10.000; AP132P); goat anti mouse HRP (1:10.000, AP124P).

### Thin-layer chromatography

Lipids associated with the total cell lysates were extracted twice with chloroform/methanol 2:1 (v:v) and chloroform/methanol/water 20:10:1 (v:v). Total lipid extracts were subjected to phase-separation according to the Folch method, with some modifications. Finally, the organic phases were subjected to alkaline methanolysis to remove glycerophospholipids (98, 99). Lipids contained in the organic and aqueous phase fractions corresponding to the same amount of cell proteins were separate by high pressure thin layer chromatography (HPTLC) using the solvent system chloroform:methanol:water 110:40:6 (v:v:v) and chloroform:methanol:CaCl_2_ 0.2%, 50:42:11 (v:v:v), respectively. Gangliosides were visualized by Ehrlich reagents, whereas the lipids contained in the organic phase were observed using anisaldehyde staining. Lipid bands were detected by digital acquisition and quantified using ImageJ software. After separation, the lipids were identified by co-migration with authentic lipid standards.

### Statistics

Data were analyzed with Graph Pad Prism version 8.0 for Macintosh (San Diego, California, USA) and expressed as the mean ± standard error of the mean (SEM). Unpaired Student t-test and one-way or two-way ANOVA followed by post-tests were used when appropriate (statistical significance: P < 0.05). Survival curves were analyzed with log-rank (Mantel–Cox) test. The number of samples and statistical tests used are indicated in the legends to each figure.

### Study approval

All experiments and procedures described in this study were conducted under an approved protocol of the Institutional Committee for the Good Animal Experimentation of the San Raffaele Scientific Institute and were reported to the Ministry of Health, as required by Italian law (IACUC #791, #1145).

## AUTHOR CONTRIBUTIONS

DS, FO conducted experiments, acquired, and analyzed data; FM, CA, SM performed and analyzed enzymatic activity assays and DEAE chromatography; VA, RDG, AB performed and analyzed morphometric and ultrastructural analysis on CNS and PNS tissues; MV and MA performed and analyzed TLC to assess ganglioside content. AB, SM, FM, MA provided reagents, intellectual input, and critical review of the manuscript; DS, FO and AG designed the research study, analyzed data, and wrote the manuscript.

## ACKNOWLEDGMENTS

We are grateful to Luigi Tiradani for LV preparation, Martina Bazzucchi for help with enzymatic assays, and all the members of Gritti’s lab for generous support and advice. Part of this work was carried out in ALEMBIC, an advanced microscopy laboratory established by IRCCS Ospedale San Raffaele and Università Vita-Salute San Raffaele. This study was funded by grants from Fondazione Telethon (Tiget Core Grant 2016-2021, project D2), National Tay-Sachs and Allied Diseases (NTSAD; 2016 grant), and Vaincre les Maladies Lysosomiales (VML, agreement 2018-4) to AG. The sponsor(s) had no role in the study design or the collection, analysis, and interpretation of data; the writing of the report; or the decision to submit the article for publication. D.S. conducted this study to fulfill his Ph.D. in Medical Biotechnologies XXXIII cycle, University of Perugia, Italy.

## REFERENCES

1. Regier, D.S., Proia, R.L., D’Azzo, A. and Tifft, C.J. (2016) The GM1 and GM2 Gangliosidoses: Natural History and Progress toward Therapy. Pediatr. Endocrinol. Rev., 13, 663–673.

2. Mahuran, D.J. (1999) Biochemical consequences of mutations causing the GM2 gangliosidoses. Biochim Biophys Acta, 1455, 105–138.

3. Cordeiro, P., Hechtman, P. and Kaplan, F. (2000) The GM2 gangliosidoses databases: allelic variation at the HEXA, HEXB, and GM2A gene loci. Genet. Med., 2, 319–27.

4. Bley, A.E., Giannikopoulos, O.A., Hayden, D., et al.,. (2011) Natural history of infantile G(M2) gangliosidosis. Pediatrics, 128, e1233–41.

5. Tsuji, D., Akeboshi, H., Matsuoka, K. Y., et al. (2011) Highly phosphomannosylated enzyme replacement therapy for GM2 gangliosidosis. Ann. Neurol., 69, 691–701.

6. Matsuoka, K., Tamura, T., Tsuji, D., et al. (2011) Therapeutic potential of intracerebroventricular replacement of modified human β-hexosaminidase B for GM2 gangliosidosis. Mol. Ther., 19, 1017–1024.

7. Baek, R.C., Kasperzyk, J.L., Platt, F.M. and Seyfried, T.N. (2008) N-butyldeoxygalactonojirimycin reduces brain ganglioside and GM2 content in neonatal Sandhoff disease mice. Neurochem. Int., 52, 1125–1133.

8. Andersson, U., Smith, D., Jeyakumar, M., et al (2004) Improved outcome of N-butyldeoxygalactonojirimycin-mediated substrate reduction therapy in a mouse model of Sandhoff disease. Neurobiol Dis, 16, 506–515.

9. Maegawa, G.H.B., Tropak, M., Buttner, J., et al. (2007) Pyrimethamine as a potential pharmacological chaperone for late-onset forms of GM2 gangliosidosis. J Biol Chem, 282, 9150– 9161.

10. Maegawa, G.H.B., Banwell, B.L., Blaser, S., et al (2009) Substrate reduction therapy in juvenile GM2 gangliosidosis. Mol. Genet. Metab., 98, 215–224.

11. Tallaksen, C.M.E. and Berg, J.E. (2009) Miglustat therapy in juvenile Sandhoff disease. J. Inherit. Metab. Dis., 32.

12. Masciullo, M., Santoro, M., Modoni, A., et al (2010) Substrate reduction therapy with miglustat in chronic GM2 gangliosidosis type Sandhoff: Results of a 3-year follow-up. J. Inherit. Metab. Dis., 33.

13. Wortmann, S.B., Lefeber, D.J., Dekomien, G., et al (2009) Substrate deprivation therapy in juvenile Sandhoff disease. J. Inherit. Metab. Dis., 32, S307–S311.

14. Clarke, J.T.R., Mahuran, D.J., Sathe, S., et al (2011) An open-label Phase I/II clinical trial of pyrimethamine for the treatment of patients affected with chronic GM2 gangliosidosis (Tay-Sachs or Sandhoff variants). Mol. Genet. Metab., 102, 6–12.

15. von Specht, B.U., Geiger, B., Arnon, R., et al (1979) Enzyme replacement in Tay-Sachs disease. Neurology, 29, 848–854.

16. Shapiro, B.E., Pastores, G.M. Gianutsos, et al (2009) Miglustat in late-onset Tay-Sachs disease: A 12-month, randomized, controlled clinical study with 24 months of extended treatment. Genet. Med., 10.1097/GIM.0b013e3181a1b5c5.

17. Sango, K., Yamanaka, S., Hoffmann, A., et al. (1995) Mouse models of Tay-Sachs and Sandhoff diseases differ in neurologic phenotype and ganglioside metabolism. Nat Genet, 11, 170–176.

18. Norflus, F., Tifft, C.J., McDonald, M.P., et al (1998) Bone marrow transplantation prolongs life span and ameliorates neurologic manifestations in Sandhoff disease mice. J. Clin. Invest., 101, 1881– 1888.

19. Wada, R., Tifft, C.J. and Proia, R.L. (2000) Microglial activation precedes acute neurodegeneration in Sandhoff disease and is suppressed by bone marrow transplantation. Proc. Natl. Acad. Sci. U. S. A., 97, 10954–10959.

20. Hoogerbrugge, P.M., Brouwer, O.F., Bordigoni, P., et al. (1995) Allogeneic bone marrow transplantation for lysosomal storage diseases. The European Group for Bone Marrow Transplantation. Lancet, 345, 1398–1402.

21. Jacobs, J.F., Willemsen, M.A., Groot-Loonen, J.J., et al (2005) Allogeneic BMT followed by substrate reduction therapy in a child with subacute Tay-Sachs disease. Bone Marrow Transpl., 36, 925– 926.

22. Cachon-Gonzalez, M.B., Wang, S.Z., Ziegler, R. C., et al (2014) Reversibility of neuropathology in Tay-Sachs-related diseases. Hum. Mol. Genet., 23, 730–748.

23. Sargeant, T.J., Wang, S., Bradley, J., et al (2011) Adeno-associated virus-mediated expression of beta-hexosaminidase prevents neuronal loss in the Sandhoff mouse brain. Hum. Mol. Genet., 20, 4371–4380.

24. McCurdy, V.J., Rockwell, H.E., Arthur, J.R., et al. (2015) Widespread correction of central nervous system disease after intracranial gene therapy in a feline model of Sandhoff disease. Gene Ther., 22, 181–189.

25. Rockwell, H.E., McCurdy, V.J., Eaton, S.C., et al. (2015) AAV-mediated gene delivery in a feline model of Sandhoff disease corrects lysosomal storage in the central nervous system. ASN Neuro, 7.

26. Bradbury, A.M., Gray-Edwards, H.L., Shirley, J.L., et al. (2015) Biomarkers for disease progression and AAV therapeutic efficacy in feline Sandhoff disease. Exp. Neurol., 263, 102–112.

27. Golebiowski, D., van der Bom, I.M.J., Kwon, C.-S., et al. (2017) Direct Intracranial Injection of AAVrh8 Encoding Monkey β-N-Acetylhexosaminidase Causes Neurotoxicity in the Primate Brain. Hum. Gene Ther., 28, 510–522.

28. Mingozzi, F. and High, K.A. (2011) Immune responses to AAV in clinical trials. Curr Gene Ther, 11, 321–330.

29. Bradbury, A.M., Cochran, J.N., McCurdy, V.J., et al. (2013) Therapeutic response in feline sandhoff disease despite immunity to intracranial gene therapy. Mol. Ther., 21, 1306–1315.

30. Hordeaux, J., Buza, E.L., Dyer, C., et al. (2020) Adeno-Associated Virus-Induced Dorsal Root Ganglion Pathology. Hum. Gene Ther., 31, 808–818.

31. Lattanzi, A., Neri, M., Maderna, C., et al (2010) Widespread enzymatic correction of CNS tissues by a single intracerebral injection of therapeutic lentiviral vector in leukodystrophy mouse models. Hum. Mol. Genet., 19, 2208–2227.

32. Lattanzi, A., Salvagno, C., Maderna, C., et al (2014) Therapeutic benefit of lentiviral-mediated neonatal intracerebral gene therapy in a mouse model of globoid cell leukodystrophy. Hum. Mol. Genet., 23, 3250–3268.

33. Palfi, S., Gurruchaga, J.M., Ralph, G.S., et al. (2014) Long-term safety and tolerability of ProSavin, a lentiviral vector-based gene therapy for Parkinson’s disease: a dose escalation, open-label, phase 1/2 trial. Lancet, 383, 1138–1146.

34. Palfi, S., Gurruchaga, J.M., Lepetit, H., et al. (2018) Long-Term Follow-Up of a Phase I/II Study of ProSavin, a Lentiviral Vector Gene Therapy for Parkinson’s Disease. Hum. Gene Ther. Clin. Dev., 29, 148–155.

35. Meneghini, V., Lattanzi, A., Tiradani, L., et al. (2016) Pervasive supply of therapeutic lysosomal enzymes in the CNS of normal and Krabbe-affected non-human primates by intracerebral lentiviral gene therapy. EMBO Mol. Med., 8, 489–510.

36. Ornaghi, F., Sala, D., Tedeschi, F., et al (2020) Novel bicistronic lentiviral vectors correct β-Hexosaminidase deficiency in neural and hematopoietic stem cells and progeny: implications for in vivo and ex vivo gene therapy of GM2 gangliosidosis. Neurobiol. Dis., 134, 104667.

37. Ogawa, Y., Sasanuma, Y., Shitara, S., et al (2019) Abnormal organization during neurodevelopment in a mouse model of Sandhoff disease. Neurosci. Res., 10.1016/j.neures.2019.07.004.

38. Ogawa, Y., Kaizu, K., Yanagi, Y., et al (2017) Abnormal differentiation of Sandhoff disease model mouse-derived multipotent stem cells toward a neural lineage. PLoS One, 10.1371/journal.pone.0178978.

39. Higami, S., Nishizawa, K., Omura, K., et al (1976) Prenatal diagnosis and fetal pathology of Tay-Sachs disease. Tohoku J Exp Med, 118, 323–330.

40. Omura, K., Higami, S., Issiki, G., et al (1973) Prenatal diagnosis of the Hurler syndrome: mucopolysaccharide pattern in amniotic fluid. Tohoku J Exp Med, 111, 87–91.

41. Higami, S., Omura, K., Nishizawa, K. Y, et al (1978) Prenatal diagnosis and fetal pathology of Niemann-Pick disease. Tohoku J Exp Med, 125, 11–17.

42. Igisu, H. and Suzuki, K. (1984) Progressive accumulation of toxic metabolite in a genetic leukodystrophy. Science (80-.)., 224, 753–755.

43. Santambrogio, S., Ricca, A., Maderna, C., et al. (2012) The galactocerebrosidase enzyme contributes to maintain a functional neurogenic niche during early post-natal CNS development. Hum. Mol. Genet., 21, 4732–4750.

44. Patil, S.A., Hb Maegawa, G. and Maegawa, G.H. (2013) Developing therapeutic approaches for metachromatic leukodystrophy. Drug Des. Devel. Ther., 7, 729–745.

45. Sandhoff, K. and Harzer, K. (2013) Gangliosides and gangliosidoses: principles of molecular and metabolic pathogenesis. J. Neurosci., 33, 10195–208.

46. Ogawa, Y., Irisa, M., Sano, T., et al (2018) Improvement in dysmyelination by the inhibition of microglial activation in a mouse model of Sandhoff disease. Neuroreport, 10.1097/WNR.0000000000001060.

47. Hasegawa, D., Tamura, S., Nakamoto, Y., et al (2013) Magnetic resonance findings of the corpus callosum in canine and feline lysosomal storage diseases. PLoS One, 8.

48. Arthur, J.R., Lee, J.P., Snyder, E.Y. and Seyfried, T.N. (2012) Therapeutic effects of stem cells and substrate reduction in juvenile Sandhoff mice. Neurochem. Res., 37, 1335–1343.

49. Lee, J.P., Jeyakumar, M., Gonzalez, R., et al. (2007) Stem cells act through multiple mechanisms to benefit mice with neurodegenerative metabolic disease. Nat. Med., 13, 439–447.

50. Jeyakumar, M., Norflus, F., Tifft, C.J., et al. (2001) Enhanced survival in Sandhoff disease mice receiving a combination of substrate deprivation therapy and bone marrow transplantation. Blood, 97, 327–329.

51. Ricca, A., Rufo, N., Ungari, S., et al. (2015) Combined gene/cell therapies provide long-term and pervasive rescue of multiple pathological symptoms in a murine model of globoid cell leukodystrophy. Hum. Mol. Genet., pii: ddv08, 3372–3389.

52. Visigalli, I., Moresco, R.M.R.M., Belloli, S., et al. (2009) Monitoring disease evolution and treatment response in lysosomal disorders by the peripheral benzodiazepine receptor ligand PK11195. Neurobiol. Dis., 34, 51–62.

53. Ornaghi, F., Sala, D., Tedeschi, F., et al (2020) Novel bicistronic lentiviral vectors correct β-Hexosaminidase deficiency in neural and hematopoietic stem cells and progeny: implications for in vivo and ex vivo gene therapy of GM2 gangliosidosis. Neurobiol. Dis., 134.

54. Capotondo, A., Milazzo, R., Salvatore, L., et al. (2012) Brain conditioning is instrumental for successful microglia reconstitution following hematopoietic stem cell transplantation. Proc. Natl. Acad. Sci. U. S. A., 109, 15018–23.

55. Ballabio, A. and Gieselmann, V. (2009) Lysosomal disorders: from storage to cellular damage. Biochim Biophys Acta, 1793, 684–696.

56. Lecommandeur, E., Cachón-González, M.B., Boddie, S., et al (2021) Decrease in myelin-associated lipids precedes neuronal loss and glial activation in the CNS of the sandhoff mouse as determined by metabolomics. Metabolites, 11, 1–16.

57. Jeyakumar, M., Thomas, R., Elliot-Smith, E., et al. (2003) Central nervous system inflammation is a hallmark of pathogenesis in mouse models of GM1 and GM2 gangliosidosis. Brain, 126, 974– 987.

58. Porter, B.F., Lewis, B.C., Edwards, J.F., et al (2011) Pathology of GM2 gangliosidosis in Jacob sheep. Vet. Pathol., 48, 807–813.

59. Baek, R.C., Martin, D.R., Cox, N.R. and Seyfried, T.N. (2009) Comparative analysis of brain lipids in mice, cats, and humans with Sandhoff disease. Lipids, 44, 197–205.

60. Cachón-González, M.B., Wang, S.Z., Ziegler, R., et al (2014) Reversibility of neuropathology in tay-sachs-related diseases. Hum. Mol. Genet., 10.1093/hmg/ddt459.

61. Haberland, C., Brunngraber, E., Witting, L. and Brown, B. (1973) The white matter in G M2 gangliosidosis. A comparative histopathological and biochemical study. Acta Neuropathol, 24, 43–55.

62. McNally, M. a, Baek, R.C., Avila, R.L., et al (2007) Peripheral nervous system manifestations in a Sandhoff disease mouse model: nerve conduction, myelin structure, lipid analysis. J. Negat. Results Biomed., 6, 8.

63. Sango, K., Yamanaka, S., Ajiki, K., et al (2002) Lysosomal storage results in impaired survival but normal neurite outgrowth in dorsal root ganglion neurones from a mouse model of Sandhoff disease. Neuropathol. Appl. Neurobiol., 28, 23–34.

64. Cachon-Gonzalez, M.B., Wang, S.Z., Lynch, A., et al. (2006) Effective gene therapy in an authentic model of Tay-Sachs-related diseases. Proc. Natl. Acad. Sci. U. S. A., 103, 10373–10378.

65. Cachon-Gonzalez, M.B., Wang, S.Z., McNair, R., et al (2012) Gene transfer corrects acute GM2 gangliosidosis--potential therapeutic contribution of perivascular enzyme flow. Mol. Ther., 20, 1489–1500.

66. Woodley, E., Osmon, K.J.L.L., Thompson, P., et al (2019) Efficacy of a Bicistronic Vector for Correction of Sandhoff Disease in a Mouse Model. 12, 47–57.

67. Lahey, H.G., Webber, C.J., Golebiowski, D., et al. (2020) Pronounced Therapeutic Benefit of a Single Bidirectional AAV Vector Administered Systemically in Sandhoff Mice. Mol. Ther., 28, 2150– 2160.

68. Niemir, N., Rouvière, L., Besse, A., et al. (2018) Intravenous administration of scAAV9-Hexb normalizes lifespan and prevents pathology in sandhoff disease mice. Hum. Mol. Genet., 27, 954–968.

69. Osmon, K.J.L., Woodley, E., Thompson, P., et al (2016) Systemic Gene Transfer of a Hexosaminidase Variant Using an scAAV9.47 Vector Corrects GM2 Gangliosidosis in Sandhoff Mice. Hum. Gene Ther., 27, 497–508.

70. Mingozzi, F. and High, K.A. (2013) Immune responses to AAV vectors: overcoming barriers to successful gene therapy. Blood, 122, 23–36.

71. Ricca, A., Rufo, N., Ungari, S., et al. (2015) Combined gene/cell therapies provide long-term and pervasive rescue of multiple pathological symptoms in a murine model of globoid cell leukodystrophy. Hum. Mol. Genet., 24, 3372–89.

72. Oya, Y., Proia, R.L., Norflus, F., et al (2000) Distribution of enzyme-bearing cells in GM2 gangliosidosis mice: regionally specific pattern of cellular infiltration following bone marrow transplantation. Acta Neuropathol, 99, 161–168.

73. Boelens, J.J., Prasad, V.K., Tolar, J., et al (2010) Current international perspectives on hematopoietic stem cell transplantation for inherited metabolic disorders. Pediatr Clin North Am, 57, 123–145.

74. Ferrari, G., Thrasher, A.J. and Aiuti, A. (2021) Gene therapy using haematopoietic stem and progenitor cells. Nat. Rev. Genet., 22, 216–234.

75. Martino, S., Marconi, P., Tancini, B., et al. (2005) A direct gene transfer strategy via brain internal capsule reverses the biochemical defect in Tay-Sachs disease. Hum Mol Genet, 14, 2113–2123.

76. Sevenich, L. (2018) Brain-resident microglia and blood-borne macrophages orchestrate central nervous system inflammation in neurodegenerative disorders and brain cancer. Front. Immunol., 9, 1–16.

77. Liddelow, S.A. and Barres, B.A. (2017) Reactive Astrocytes: Production, Function, and Therapeutic Potential. Immunity, 46, 957–967.

78. Tremblay, M. and Sierra, A. Microglia in Health and Disease (2014) Springer New York, (2014), 1–486

79. Cachon-Gonzalez, M.B., Zaccariotto, E. and Cox, T.M. (2018) Genetics and Therapies for GM2 Gangliosidosis. Curr. Gene Ther., 18, 68–89.

80. Wada, R., Tifft, C.J. and Proia, R.L. (2000) Microglial activation precedes acute neurodegeneration in Sandhoff disease and is suppressed by bone marrow transplantation. Proc. Natl. Acad. Sci. U. S. A., 97, 10954–10959.

81. Huang, J.Q., Trasler, J.M., Igdoura, S., et al (1997) Apoptotic cell death in mouse models of G(M2) gangliosidosis and observations on human Tay-Sachs and Sandhoff diseases. Hum. Mol. Genet., 6, 1879–1885.

82. Alonso-Pérez, J., Casasús, A., Gimenez-Muñoz,Á., et al (2021) Late onset Sandhoff disease presenting with lower motor neuron disease and stuttering. Neuromuscul. Disord., 10.1016/j.nmd.2021.04.011.

83. Hölzer, H.T., Boschann, F., Hennermann, J.B., et al (2021) Cerebellar atrophy on top of motor neuron compromise as indicator of late-onset GM2 gangliosidosis. J. Neurol., 268, 2259–2262.

84. Rowe, O.E., Rangaprakash, D., Weerasekera, A., et al (2021) Magnetic resonance imaging and spectroscopy in late-onset GM2-gangliosidosis. Mol. Genet. Metab., 133, 386–396.

85. Masingue, M., Dufour, L., Lenglet, T., et al. (2020) Natural History of Adult Patients with GM2 Gangliosidosis. Ann. Neurol., 87, 609–617.

86. Wessels, M.E., Holmes, J.P., Jeffrey, M., et al (2014) GM2 gangliosidosis in British Jacob sheep. J. Comp. Pathol., 150, 253–257.

87. Liu, Y., Hoffmann, A., Grinberg, A., et al (1997) Mouse model of GM2 activator deficiency manifests cerebellar pathology and motor impairment. Proc. Natl. Acad. Sci. U. S. A., 94, 8138– 43.

88. Kroll, R.A., Pagel, M.A., Roman-Goldstein, S., et al (1995) White matter changes associated with feline GM2 gangliosidosis (Sandhoff disease): correlation of MR findings with pathologic and ultrastructural abnormalities. AJNR Am J Neuroradiol, 16, 1219–1226.

89. Ito, D., Ishikawa, C., Jeffery, N.D., et al (2018) Two-Year Follow-Up Magnetic Resonance Imaging and Spectroscopy Findings and Cerebrospinal Fluid Analysis of a Dog with Sandhoff’s Disease. J. Vet. Intern. Med., 10.1111/jvim.15041.

90. Fumagalli, F., Calbi, V., Sessa, M., et al. (2020) Lentiviral hematopoietic stem and progenitor cell gene therapy (HSPC-GT) for metachromatic leukodystrophy (MLD): Clinical outcomes from 33 patients. Mol. Genet. Metab., 129(2), S59.

91. Sessa, M., Lorioli, L., Fumagalli, F., et al. (2016) Lentiviral haemopoietic stem-cell gene therapy in early-onset metachromatic leukodystrophy: an ad-hoc analysis of a non-randomised, open-label, phase 1/2 trial. Lancet, 388, 476–487.

92. Gentner, B., Tucci, F., Galimberti, S., et al. (2021) Hematopoietic Stem-and Progenitor-Cell Gene Therapy for Hurler Syndrome. N. Engl. J. Med., 385, 1929–1940.

93. Capotondo, A., Milazzo, R., Garcia-Manteiga, J.M. C, et al (2017) Intracerebroventricular delivery of hematopoietic progenitors results in rapid and robust engraftment of microglia-like cells. Sci. Adv., 3.

94. Martino, S., Di Girolamo, I., Cavazzin, C., et al. (2009) Neural precursor cell cultures from GM2 gangliosidosis animal models recapitulate the biochemical and molecular hallmarks of the brain pathology. J. Neurochem., 109, 135–147.

95. Schindelin, J., Arganda-Carreras, I., Frise, E. K, et al. (2012) Fiji: An open-source platform for biological-image analysis. Nat. Methods, 9, 676–682.

96. Wrabetz, L., Feltri, M.L., Quattrini, A., et al. (2000) P(0) glycoprotein overexpression causes congenital hypomyelination of peripheral nerves. J. Cell Biol., 148, 1021–1034.

97. Noseda, R., Guerrero-Valero, M., Alberizzi, V., et al. (2016) Kif13b Regulates PNS and CNS Myelination through the Dlg1 Scaffold. PLoS Biol., 14, 1–26.

98. Samarani, M., Loberto, N., Soldà, G., et al. (2018) A lysosome–plasma membrane–sphingolipid axis linking lysosomal storage to cell growth arrest. FASEB J., 10.1096/fj.201701512RR.

99. Valsecchi, M., Aureli, M., Mauri, L., et al. (2010) Sphingolipidomics of A2780 human ovarian carcinoma cells treated with synthetic retinoids. J. Lipid Res., 10.1194/jlr.M004010.

