## Supplementary figures_legends_table 1 for "Therapeutic advantages of combined gene/cell therapy strategies in a murine model of GM2 gangliosidosis"

### Supplementary material

- *Supplementary figures and legends*
- *Legends to Supplementary Videos*
- *Supplementary Table 1*

***Supplementary figures and legends***

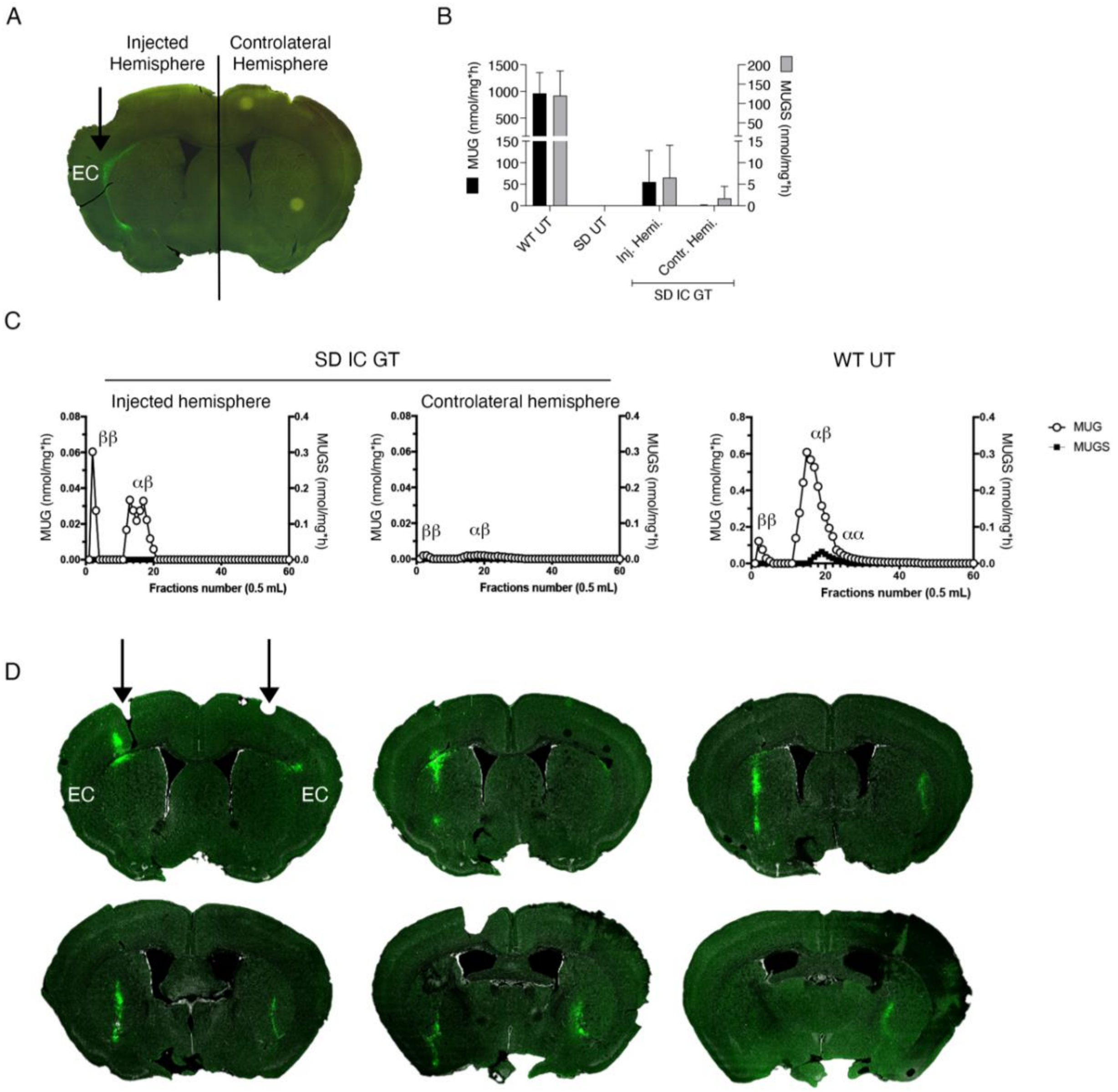


**Supplementary Figure 1. Rescue of Hex activity in SD mice following IC GT. (A)** Representative IF picture showing the distribution of GFP+ cells in coronal brain section of SD mice 30 d after neonatal monolateral injection of LV. GFP vector in white matter external capsule (EC); direct fluorescence, green. **(B)** Enzymatic activity measured by the degradation of the artificial substrates MUG (left Y-axis; nmol/mg/h) and MUGS (right Y axis; nmol/mg/h) in the Injected (Inj.) and contralateral (Contra) hemispheres (Hemi) of SD mice and UT WT and SD controls 30 d after neonatal injection (LV.mA+LV.mB). Data represent the mean ± SEM; n = 3 mice/group. Unpaired t test, Injected vs. contralateral hemispheres (MUG, MUGS), p>0.05. **(C)** Chromatographic analysis of injected and contralateral hemisphere of SD mice 30 d after unilateral injection of LV.hexa+LV.hexb showed reconstitution of Hex isoforms: αβ, HexA; ββ, HexB; αα, HexS. **(D)** Representative IF pictures showing the distribution of GFP+ cells in coronal brain sections of SD mice 60 d after neonatal bilateral injection of LV. GFP vector in white matter external capsule (EC) (arrows). Direct fluorescence, green; nuclei are counterstained with Hoechst, white.

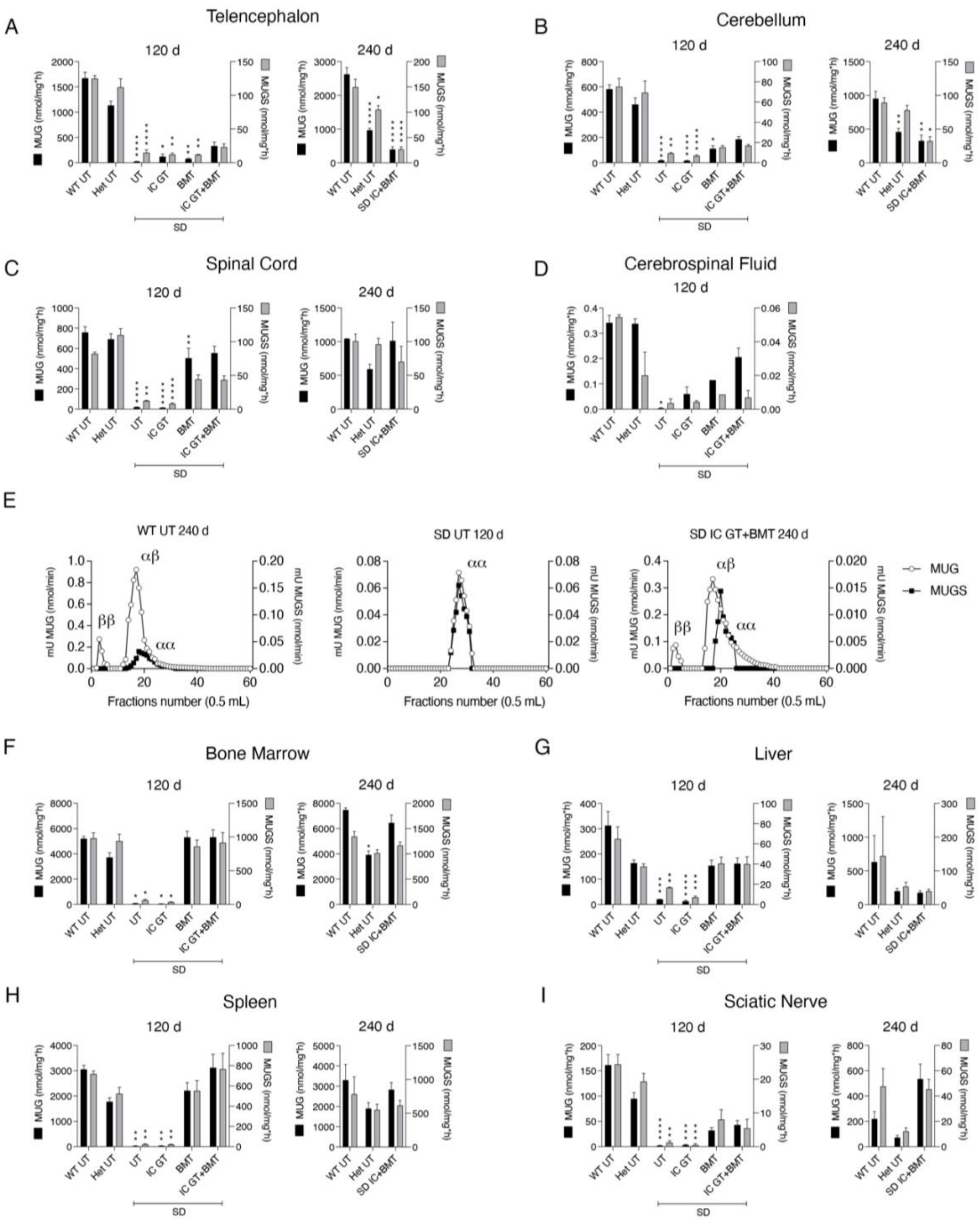


**Supplementary Figure 2. Reconstitution of Hex activity in CNS, PNS, and peripheral organs of treated SD mice.** Enzymatic activity measured as the degradation of artificial substrates MUG (left Y axis; nmol/mg/h) and MUGS (right Y axis; nmol/mg/h) in telencephalon **(A)**, cerebellum (**B**), spinal cord **(C)**, cerebrospinal fluid **(D)**, bone marrow **(F)**, liver **(G)**, spleen **(H),** and sciatic nerve **(I)** of SD treated mice (IC GT, BMT, and IC GT+BMT) and untreated (UT) controls (WT, Het, and SD) at 120 d and 240 d. Data are expressed as mean ± SEM; n = 2-9 mice/group. One-way ANOVA followed by Kruskal–Wallis multiple comparison test, *p < 0.05, **p < 0.01, ***p < 0.001, ****p < 0.0001 vs corresponding region in WT UT mice. **(E)** Diethylaminoethyl cellulose chromatography showing presence of αβ (HexA), ββ (HexB), and αα (HexS) isoforms in the telencephalon of IC GT+BMT-treated SD mouse at 240 d and age-matched UT WT (240 d) and SD (120 d) controls. Enzymatic activity (expressed as nmol/min; mU MUG on the left and mU MUGS on the right) and fraction number (0.5 ml) are plotted on the y and x axes, respectively.

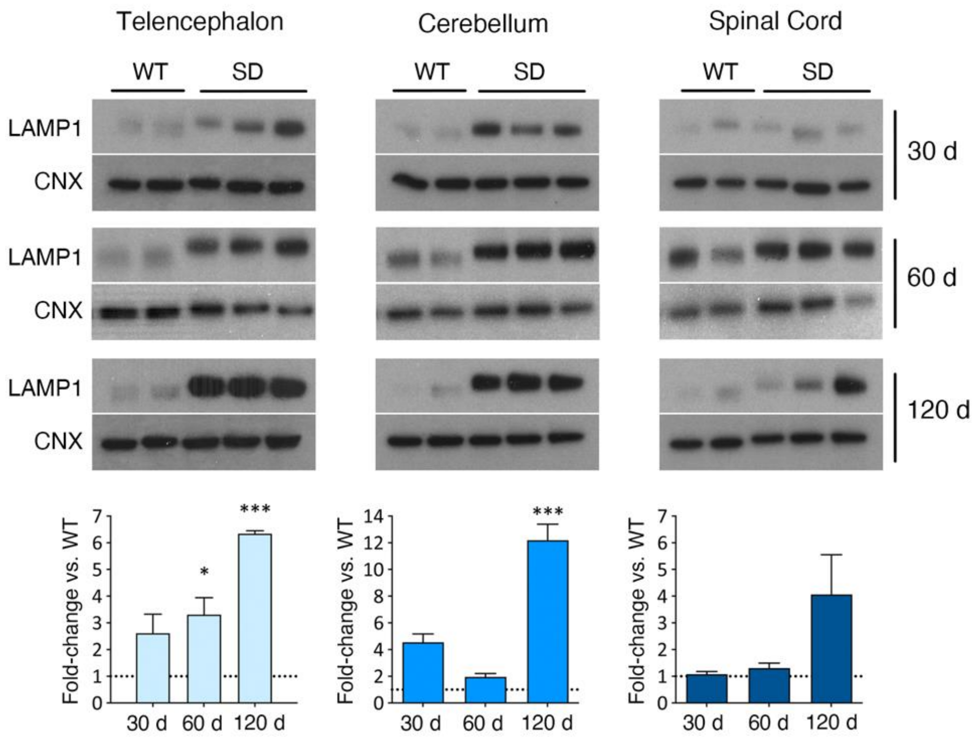


**Supplementary Figure 3. LAMP1 protein expression in SD mice at different stages of disease.** Western blot analysis and relative quantification showing LAMP1 in the TEL, CB, and SC of UT SD and WT mice at 30 d, 60 d, and 120 d of age. Data are expressed as fold change vs WT (set as 1) after normalization to calnexin (CNX). Data represent mean ± SEM; n = 2-3 mice/group. One-way ANOVA followed by Kruskal–Wallis multiple comparison test, *p < 0.05, ***p < 0.001 vs WT.

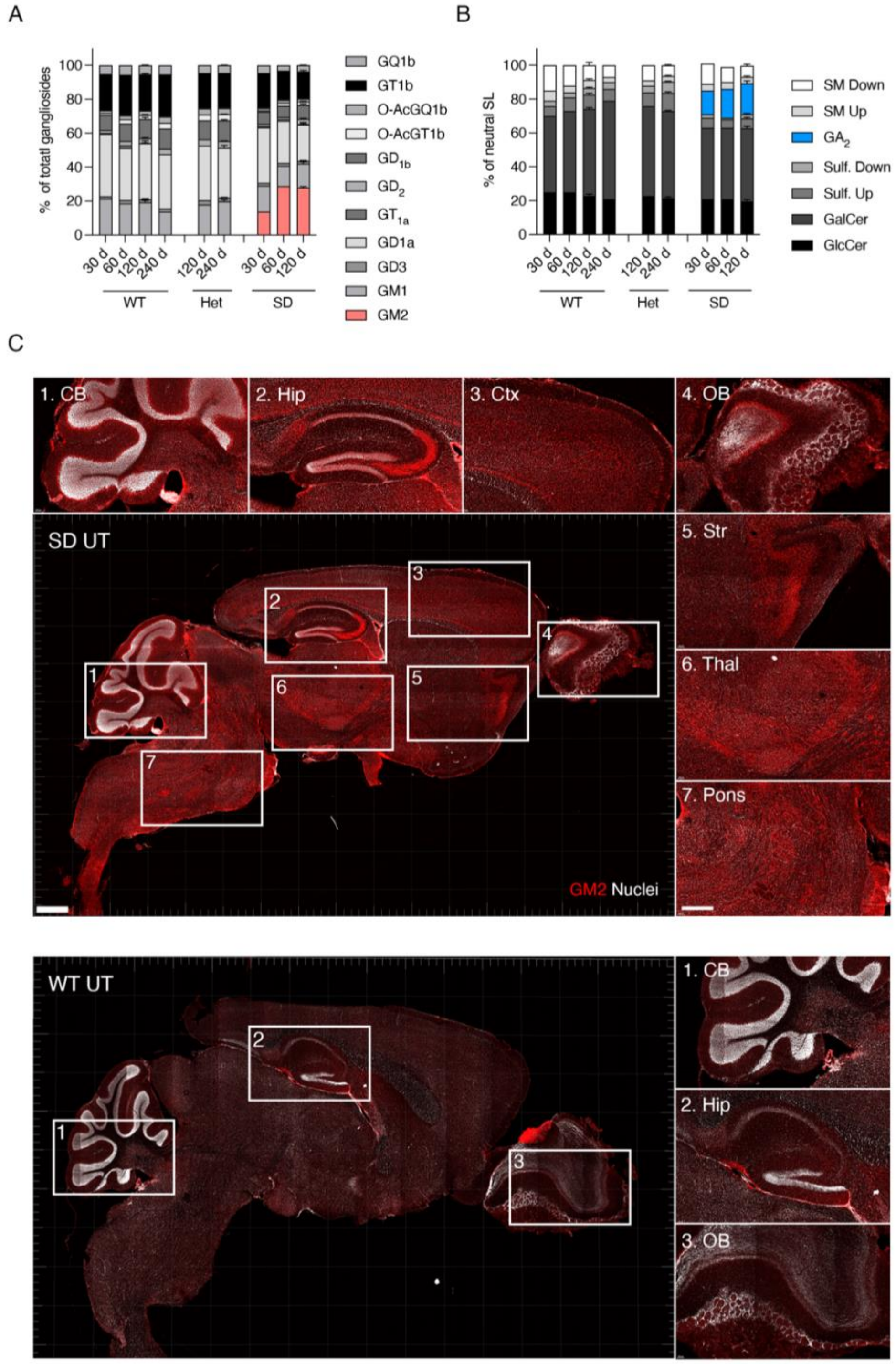


**Supplementary Figure 4. Time-course analyses of GM2 storage in untreated SD mice**. Quantification of aqueous phase **(A)** and organic phase **(B)** of total lipids obtained from brain tissue (half hemisphere) of UT WT, Het, and SD mice at 30 d, 60 d, and 120 d of age. Data are expressed as percentage of total gangliosides (aqueous phase) and percentage of total neutral sphingolipids (organic phase). Data represent mean ± SEM; n = 1-3 mice/group. GA2 and GM2 ganglioside contents were undetectable in WT and Het tissues, while they represented 28.7 ± 4.2% and 19.5 ± 4.5% of total gangliosides and total neutral sphingolipids, respectively, in SD tissues**.** All other lipid classes were similarly represented in SD, WT, and Het tissues. **(C)** Representative IF images showing GM2 storage (red; anti-GM2 antibody) in whole-brain sections of UT SD and WT mice at 120 d. Boxes highlight brain areas that are shown at higher magnification: (1) cerebellum (CB), (2) hippocampus (Hip), (3) cortex (Ctx), (4) olfactory bulb (OB), (5) striatum (Str), (6) thalamus (Thal), (7) pons, for SD; (1) cerebellum (CB), (2) hippocampus (Hip), (3) olfactory bulb (OB), for WT. GM2 is in red and nuclei are counterstained with Hoechst (white). Scale bar, 500 µm.

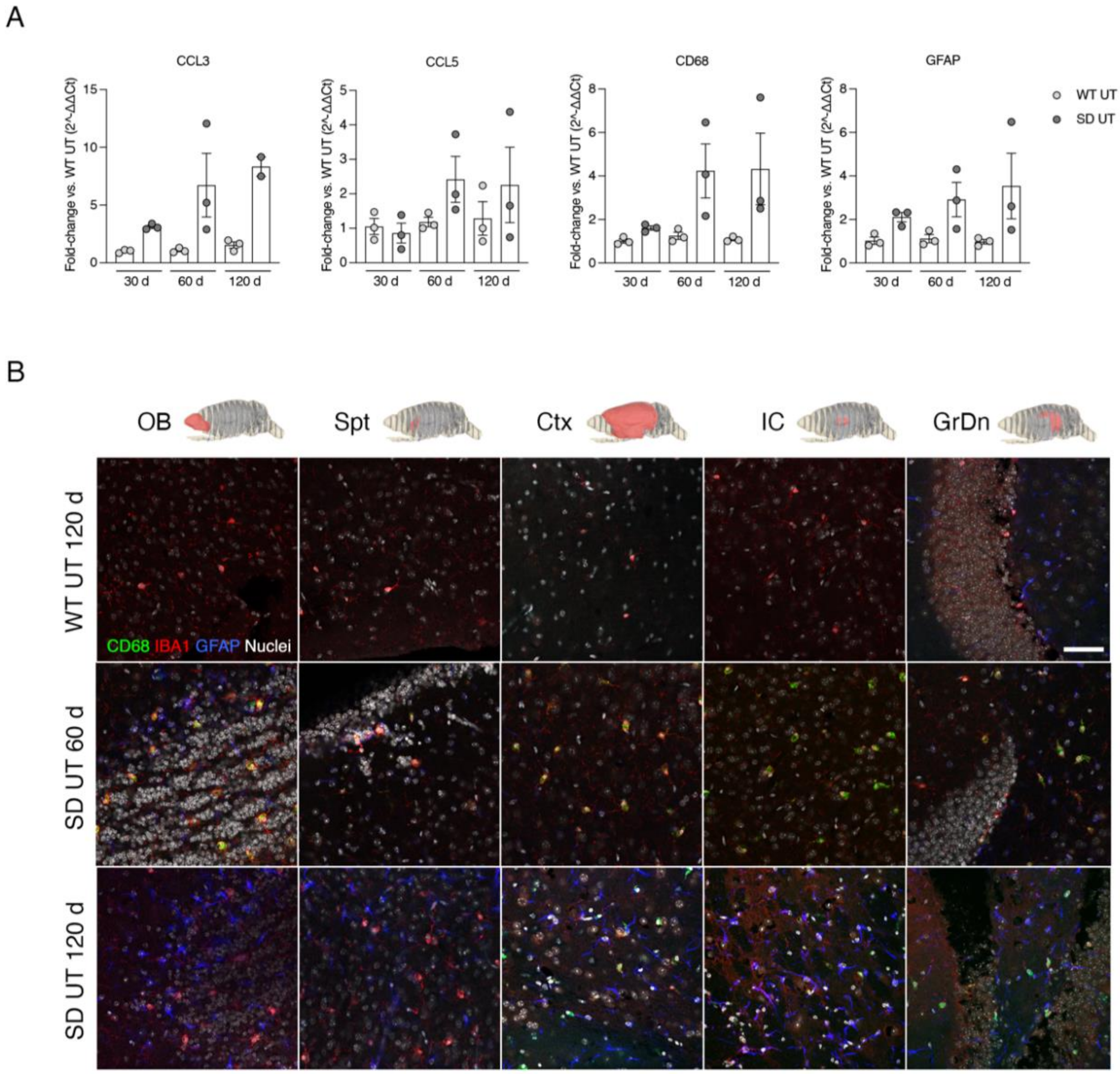


**Supplementary Figure 5. Upregulation of neuroinflammatory markers in untreated SD mice . (A)** Relative mRNA expression levels of CCL3, CCL5, CD68, and GFAP in the brain of untreated SD mice at 30 d, 60 d, and 120 d of age. Data are expressed as fold-change vs WT levels (set as 1) after normalization to GAPDH expression. Data represent mean ± SEM; n = 2-3 mice/group. One-way ANOVA followed by Sidak's multiple comparison test; SD vs age-matched WT (all genes), $p > 0.05$**. (B)** Representative confocal immunofluorescence images showing the increased expression of macrophages (CD68, green), microglia (Iba1, red), and astroglia (GFAP, blue) in the olfactory bulb (OB), septum pellucidum (Spt), cortex (Ctx), internal capsule (IC), and dentate gyrus of the hippocampus (GrDn) of untreated (UT) WT (120 d) and SD mice (60 d and 120 d). Nuclei counterstained with Hoechst (white). Scale bar, 50 µm.

**Legends to Supplementary Videos**

**Supplementary Video 1. SD UT vs SD BMT at 120 d**
Untreated SD mouse at the terminal phase of the disease (120 d) was ataxic and unable to move due to immobilization of hind limbs. SD mouse treated with BMT was still able to move but started to display signs of degeneration, e.g., bradykinesia and rigidity (spasticity) of hind limbs

**Supplementary Video 2. SD UT vs SD IC GT+BMT at 120 d**
Untreated SD mouse at the terminal phase of the disease (120 d) was ataxic and unable to move due to immobilization of hind limbs. SD mouse treated with IC GT +BMT was in better condition and maintained good mobility at 120 d.

**Supplementary Video 3. WT IC GT+BMT vs SD IC GT+BMT at 240 d**
The combination-treated SD mouse at 240 d of age (light grey hair) displayed only mild tremor and ataxia, maintained the ability to walk and feed independently, and showed explorative activity similar to the combination-treated WT littermate (dark grey hair) at timepoints far beyond the median lifespan of untreated affected SD controls or mice treated with BMT or IC GT alone.

**Supplementary Table 1**. **Evaluation of peripheral blood composition of SD and WT mice.**

Hemocytometer values of untreated (UT) WT and SD mice, tgGFP donor mice, BMT-, and IC GT+BMT-treated mice 1-month post-transplantation (90 d of age) showing subcellular composition of peripheral blood (absolute number and percentage on total no. of cells). Data are expressed as mean ± SD. RBC, red blood cells; HGB, hemoglobin; HCT, hematocrit; MCV, mean corpuscular volume; MCH, mean corpuscolar hemoglobin; MCHC, Mean Corpuscular Hemoglobin Concentration; RET, reticulocytes; PLT, platelets; WBC, white blood cells; NEUT, neutrophils; LYMPH, lymphocytes; MONO, monocytes; EO, eosinophils; BASO, basophils. #, absolute numbers.

| | RBC (M/μL) | HGB (g/dL) | HCT (%) | MCV (fL) | MCH (pg) | MCHC (g/dL) | RET# (K/μL) | RET (%) | PLT (K/μL) |
|---|---|---|---|---|---|---|---|---|---|
| **WT UT n = 7** | 10.8 ± 0.5 | 15.5 ± 0.7 | 51.5 ± 2.7 | 47.8 ± 2.2 | 14.4 ± 0.5 | 30.1 ± 0.8 | 399.0 ± 34.9 | 3.7 ± 0.3 | 1273.9 ± 297.9 |
| **tgGFP n = 11** | 10.6 ± 0.8 | 15.3 ± 0.9 | 50.2 ± 3.6 | 47.3 ± 2.2 | 14.4 ± 0.6 | 30.4 ± 0.8 | 420.7 ± 237.0 | 4.1 ± 2.7 | 1221.4 ± 320.7 |
| **SD UT n = 5** | 10.4 ± 0.2 | 15.9 ± 0.3 | 52.1 ± 0.8 | 50.3 ± 0.5 | 15.3 ± 0.3 | 30.5 ± 0.4 | 353.4 ± 98.6 | 3.4 ± 1.0 | 880.2 ± 45.4 |
| **SD BMT n = 12** | 9.8 ± 2.4 | 14.0 ± 3.3 | 45.9 ± 11.7 | 46.7 ± 2.3 | 14.3 ± 0.5 | 30.7 ± 1.5 | 312.9 ± 80.6 | 3.0 ± 0.8 | 877.1 ± 307.5 |
| **WT BMT n = 8** | 10.4 ± 1.3 | 14.8 ± 1.6 | 48.2 ± 5.9 | 46.3 ± 2.0 | 14.2 ± 0.5 | 30.7 ± 1.0 | 439.4 ± 102.9 | 4.3 ± 1.1 | 854.3 ± 265.5 |
| **SD IC GT + BMT n = 20** | 10.4 ± 0.4 | 15.0 ± 0.5 | 50.4 ± 2.2 | 48.4 ± 1.2 | 14.3 ± 0.3 | 29.7 ± 0.7 | 327.1 ± 58.4 | 3.1 ± 0.6 | 939.6 ± 250.0 |

| | WBC (K/μL) | NEUT# (K/μL) | LYMPH# (K/μL) | MONO# (K/μL) | EO# (K/μL) | BASO# (K/μL) | NEUT (%) | LYMPH (%) | MONO (%) | EO (%) | BASO (%) |
|---|---|---|---|---|---|---|---|---|---|---|---|
| **WT UT n = 7** | 9.4 ± 2.2 | 5.2 ± 3.7 | 3.8 ± 1.7 | 0.1 ± 0.2 | 0.4 ± 0.3 | 0.0 ± 0.0 | 49.6 ± 29.9 | 44.9 ± 28.6 | 1.6 ± 2.9 | 3.8 ± 2.6 | 0.1 ± 0.1 |
| **tgGFP n = 11** | 10.4 ± 2.7 | 5.1 ± 3.0 | 4.9 ± 2.7 | 0.1 ± 0.1 | 0.4 ± 0.1 | 0.0 ± 0.0 | 47.4 ± 22.1 | 48.0 ± 21.7 | 0.6 ± 0.6 | 3.8 ± 1.5 | 0.1 ± 0.1 |
| **SD UT n = 5** | 11.3 ± 2.2 | 5.5 ± 2.6 | 5.6 ± 1.4 | 0.0 ± 0.0 | 0.2 ± 0.1 | 0.0 ± 0.0 | 46.6 ± 15.9 | 51.0 ± 15.9 | 0.2 ± 0.1 | 2.2 ± 0.5 | 0.0 ± 0.1 |
| **SD BMT n = 12** | 14.5 ± 5.1 | 6.6 ± 2.0 | 7.1 ± 4.7 | 0.2 ± 0.3 | 0.6 ± 0.2 | 0.0 ± 0.0 | 48.8 ± 17.2 | 45.9 ± 15.9 | 1.2 ± 1.4 | 4.1 ± 0.9 | 0.1 ± 0.0 |
| **WT BMT n = 8** | 13.3 ± 4.6 | 7.5 ± 4.1 | 5.3 ± 2.0 | 0.1 ± 0.0 | 0.5 ± 0.4 | 0.0 ± 0.0 | 53.1 ± 15.6 | 42.8 ± 16.9 | 0.5 ± 0.3 | 3.5 ± 1.7 | 0.1 ± 0.1 |
| **SD IC GT + BMT n = 20** | 16.3 ± 4.0 | 6.6 ± 2.6 | 9.0 ± 4.0 | 0.2 ± 0.2 | 0.5 ± 0.2 | 0.0 ± 0.0 | 41.3 ± 15.6 | 54.4 ± 15.4 | 1.0 ± 0.8 | 3.2 ± 0.9 | 0.1 ± 0.1 |
| **WT IC GT + BMT n = 5** | 13.5 ± 2.8 | 5.9 ± 3.3 | 7.1 ± 1.9 | 0.1 ± 0.1 | 0.4 ± 0.1 | 0.0 ± 0.0 | 42.1 ± 16.1 | 54.3 ± 16.5 | 0.8 ± 0.9 | 2.9 ± 1.0 | 0.1 ± 0.1 |
